# Anipose: a toolkit for robust markerless 3D pose estimation

**DOI:** 10.1101/2020.05.26.117325

**Authors:** Pierre Karashchuk, Katie L. Rupp, Evyn S. Dickinson, Sarah Walling-Bell, Elischa Sanders, Eiman Azim, Bingni W. Brunton, John C. Tuthill

## Abstract

Quantifying movement is critical for understanding animal behavior. Advances in computer vision now enable markerless tracking from 2D video, but most animals live and move in 3D. Here, we introduce Anipose, a Python toolkit for robust markerless 3D pose estimation. Anipose is built on the popular 2D tracking method DeepLabCut, so users can easily expand their existing experimental setups to obtain accurate 3D tracking. It consists of four components: (1) a 3D calibration module, (2) filters to resolve 2D tracking errors, (3) a triangulation module that integrates temporal and spatial regularization, and (4) a pipeline to structure processing of large numbers of videos. We evaluate Anipose on four datasets: a moving calibration board, fruit flies walking on a treadmill, mice reaching for a pellet, and humans performing various actions. By analyzing 3D leg kinematics tracked with Anipose, we identify a key role for joint rotation in motor control of fly walking. We believe this open-source software and accompanying tutorials (anipose.org) will facilitate the analysis of 3D animal behavior and the biology that underlies it.

## 1 Introduction

Tracking body kinematics is key to answering questions in many scientific disciplines. For example, neuroscientists quantify animal movement to relate it to brain dynamics [1, 2], biomechanists quantify the movement of specific body structures to understand their mechanical properties [3, 4], social scientists quantify the motion of multiple individuals to understand their interactions [5, 6], and rehabilitation scientists quantify body movement to diagnose and treat disorders [7, 8, 9]. In all of these disciplines, achieving rapid and accurate quantification of animal pose is a major bottleneck to scientific progress.

While it is possible for human observers to recognize body movements, scoring behaviors by eye is laborious and often fails to detect differences in the rapid, fine-scale movements that characterize many behaviors. Methods for automated tracking of body kinematics from video have existed for many years, but they typically rely on the addition of markers to identify and disambiguate body parts. Although such methods can achieve very precise pose estimation [10], the use of markers is often impractical, particularly when studying natural behaviors in complex environments, tracking multiple body parts, or studying small animals. Thus, there is a pressing need for methods that perform automated, markerless tracking of body kinematics.

Recent advances in computer vision and machine learning have dramatically improved the speed and accuracy of markerless body pose estimation [1]. There are now a number of tools that apply these methods to track animal movement from 2D videos, such as DeepLabCut [11], SLEAP [12], DeepPoseKit [13], among others [14, 15, 16]. These software packages allow users to label keypoints, train convolutional neural networks, and apply them to identify keypoints from new data; several toolkits also include auxiliary tools, such as visualizing and filtering the tracked keypoints. Among them, DeepLabCut is the most widely used [17].

While tracking of animal movement from 2D video is useful for monitoring specific body parts, full body pose estimation and measurement of complex or subtle behaviors require tracking in three dimensions. Multiple tools have emerged for 3D tracking and body pose estimation, including DANNCE [18], FreiPose [19], DeepFly3D [20], and OpenMonkeyStudio [21]. However, these tools use fundamentally distinct network architectures, workflows, and user interfaces from popular 2D tracking methods. Out of the existing 2D tracking tools, only DeepLabCut [22] supports triangulation with up to 2 cameras. However, three or more cameras are often required to resolve pose ambiguities, such as when one body part occludes another. Thus, there is a need for additional tools that allow users to extend their existing 2D tracking setups to achieve robust 3D pose estimation while preserving their established workflows.

Here, we introduce Anipose (a portmanteau of “animal” and “pose”), a new toolkit to quantify 3D body kinematics by integrating DeepLabCut tracking from multiple camera views. Anipose consists of a robust calibration module, filters to further refine 2D and 3D tracking, and an interface to visualize and annotate tracked videos (example here). These features allow users to analyze 3D animal movement by extracting behavior and kinematics from videos in a unified software framework. Below, we demonstrate the value of 3D tracking with Anipose for analysis of mice, fly, and human body kinematics (Figure 1). Applying 3D tracking to estimate joint angles of walking *Drosophila*, we find that flies move their middle legs primarily by rotating their coxa and femur, whereas the front and rear legs are driven primarily by femur-tibia flexion. We then show how Anipose can be used to quantify differences between successful and unsuccessful trajectories in a mouse reaching task. Finally, we visualize how specific leg joint angles map onto a manifold of human walking.

**Figure 1:**
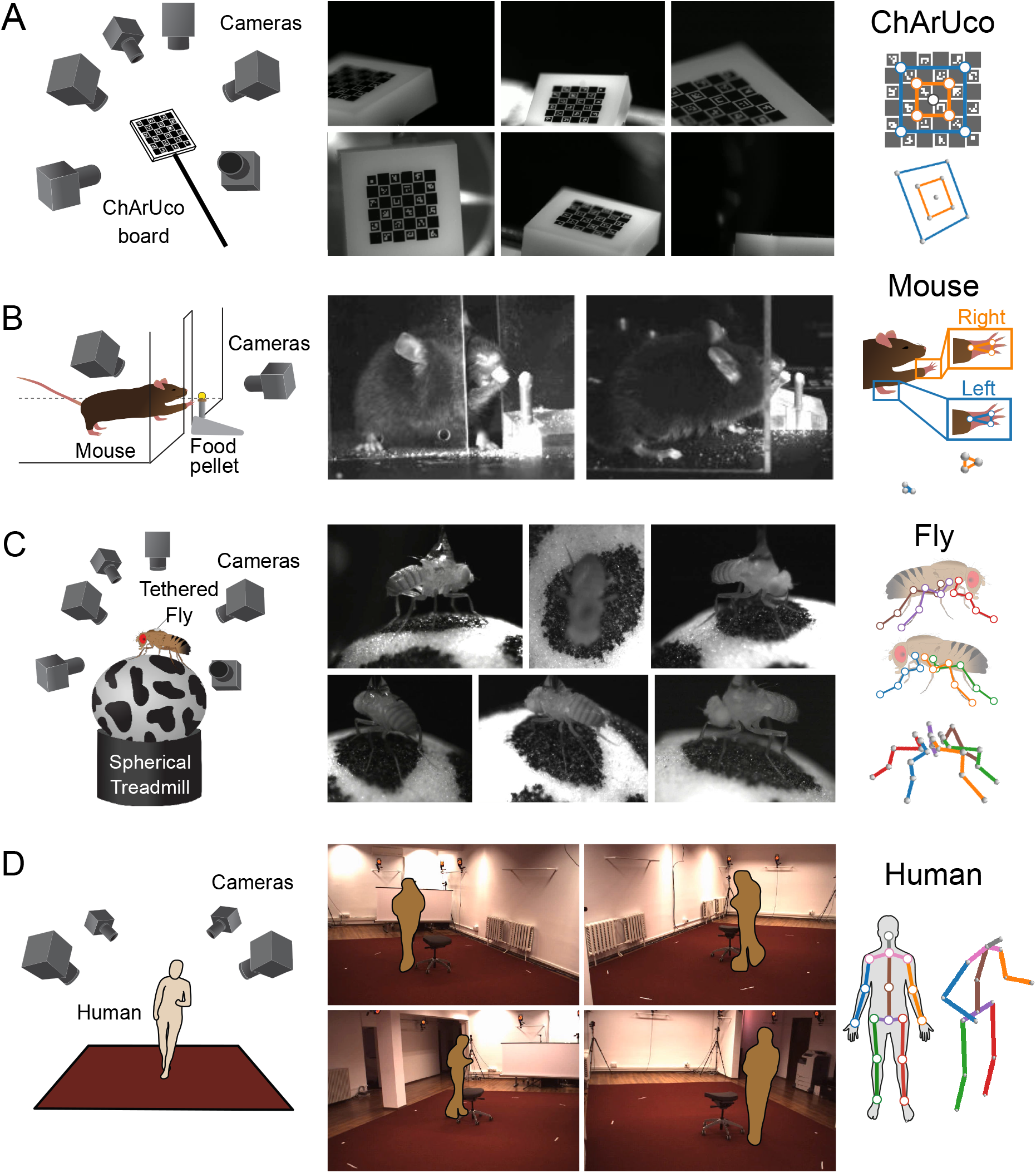
Four experimental datasets were used for evaluating 3D calibration and tracking with Anipose. (A) To evaluate tracking errors, a 2 × 2mm precision manufactured ChArUco board was simultaneously filmed from 6 cameras focused on the same point in space. We manually annotated and tracked 9 keypoints on the ChArUco board, a subset of the points that can be detected automatically with OpenCV. (B) Adult mice were trained to reach for food pellets through an opening in a clear acrylic box. After training, reach attempts were captured from 2 cameras. To quantify reach kinematics, we labeled and tracked 3 keypoints on each hand. (C) Fruit flies were tethered and positioned on a spherical treadmill, where they were able to walk, groom, etc. Fly behavior was filmed from 6 cameras evenly distributed around the treadmill. We labeled and tracked 5 keypoints on each of the 6 legs, one keypoint for each of the major leg joints. (D) As part of the Human 3.6M dataset, professional actors performing a range of actions were filmed from 4 cameras. We tracked 17 joints on each human, covering the major joints of the human body. (The images were anonymized to comply with bioRxiv guidelines.)

We designed Anipose to make 3D tracking accessible for a broad community of scientists. Because it is built on DeepLabCut, Anipose allows users to easily upgrade from 2D to 3D tracking, as well as take advantage of the DeepLabCut community, documentation, and continued support. To help new users get started, we provide in-depth tutorials and documentation at http://anipose.org. The release of Anipose as free and open-source Python software facilitates adoption, promotes ongoing contributions by community developers, and supports open science.

## 2 Results

We implement 3D tracking in a series of steps: estimation of calibration parameters from calibration videos, detection and refinement of 2D joint keypoints, triangulation and refinement of keypoints to obtain 3D joint positions, and computation of joint angles (Figure 2). In addition to the processing pipeline, the key innovations of Anipose are a robust 3D calibration module, spatiotemporal filters that refine pose estimation in both 2D and 3D, and a visualization incorporating videos, tracked keypoints, and behavioral annotations in one interface. We evaluated the calibration and triangulation modules without filters by testing their ability to accurately estimate lengths and angles of a calibration board with known dimensions (Figure 1A) and to track the hand of a mouse reaching for a food pellet (Figure 1B). We then evaluated how filtering improves estimation in 3D of position and time derivative of walking flies (Figure 1C) and humans (Figure 1D). Representative examples of tracking from each dataset are shown in Video 1.

**Figure 2:**
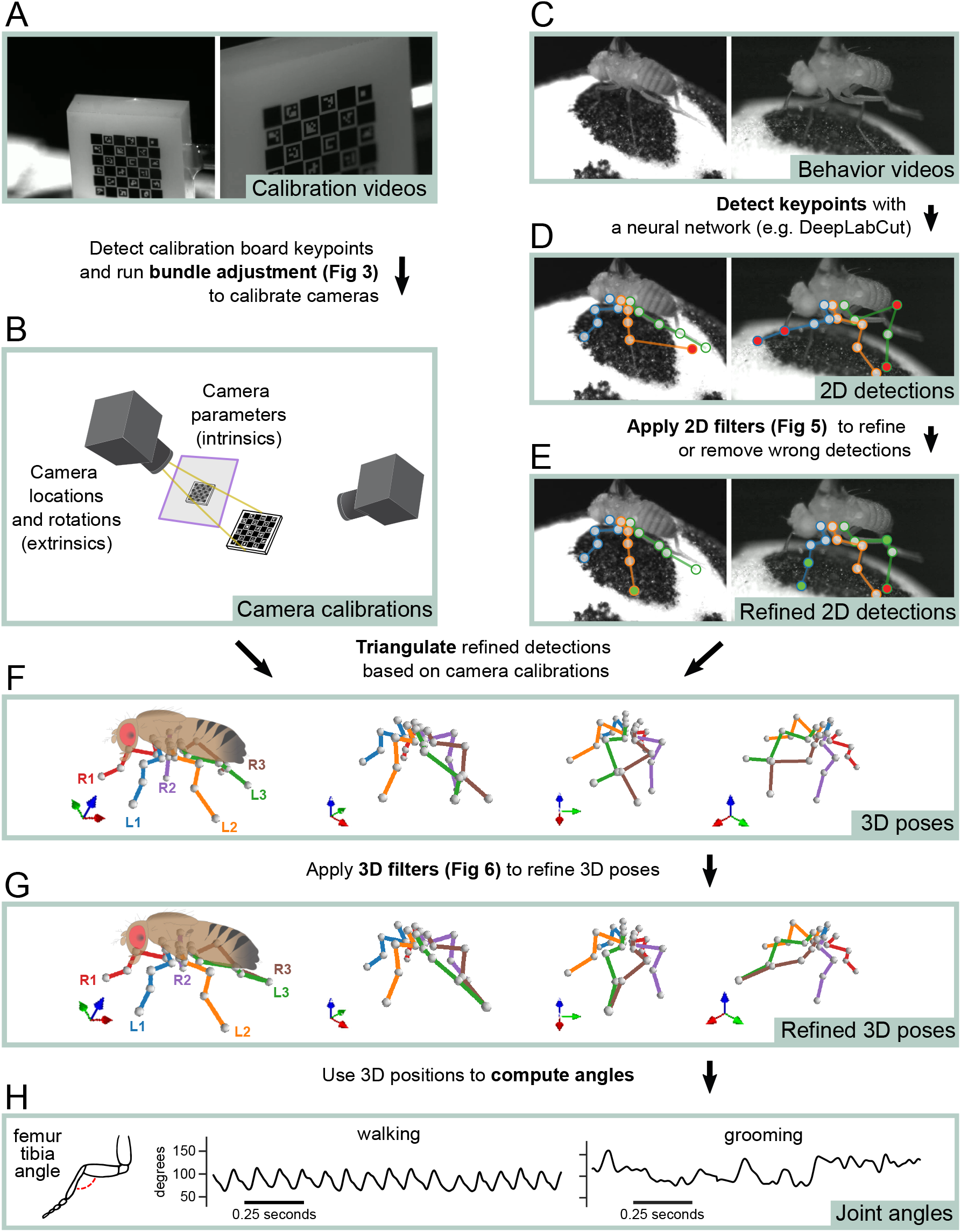
Overview of the Anipose 3D tracking pipeline. (A) The user collects simultaneous video of a calibration board from multiple cameras. (B) Calibration board keypoints are detected from calibration videos and processed to calculate intrinsic and extrinsic parameters for each camera using iterative bundle adjustment (Figure 3). (C) With the same hardware setup as in A, the user collects behavior videos. (D) Behavior videos are processed by a neural network (e.g., DeepLabCut) to detect 2D keypoints. (E) 2D keypoints are refined with 2D filters to obtain refined 2D detections (Figure 6). (F) The filtered 2D keypoints are triangulated to estimate 3D poses. (G) The estimated 3D poses are passed through an additional spatiotemporal filtering step to obtain refined 3D poses (Figure 7). (H) Joint angles are extracted from the refined 3D poses for further analysis.

### 2.1 Robust calibration of multiple camera views

An essential step in accurate 3D pose estimation is precise camera calibration, which determines the relative location and parameters of each camera (i.e., the focal length and distortions). We implemented an automated procedure that calibrates the cameras from simultaneously acquired videos of a standard calibration board (e.g., checkerboard or ChArUco board) moved by hand through the cameras’ fields of view (Figure 2A). We recommend the ChArUco board because its keypoints may be detected even with partial occlusion and its rotation can be determined uniquely from multiple views. The pipeline starts by detecting keypoints on the calibration board automatically using OpenCV [23], based on the board’s geometric regularities (e.g., checkerboard grid pattern, specific black and white markers). These board detections are used first to initialize camera calibration parameters from arbitrary positions through a greedy algorithm which adds edges between cameras one by one until it reaches a fully connected tree (Figure 3A).

**Figure 3:**
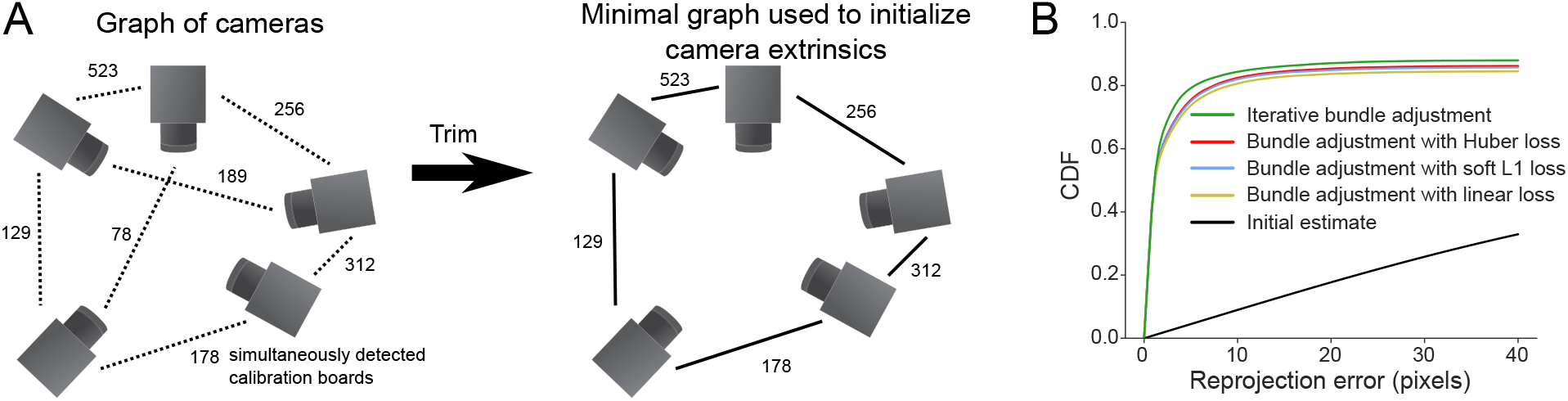
(A) Illustration of the camera parameter initialization procedure. We build a graph with each camera as a node and edge weights computed by the number of frames the calibration board is simultaneously detected by pairs of cameras. To initialize the camera calibration, we trim this graph to be a minimal, fully connected tree using a greedy approach. (B) On calibration videos from the fly dataset, bundle adjustment improves the initial calibration estimate, as measured by a reduction in reprojection error.

Although some tracking tools (e.g., [14, 18]) stop at the initial estimate of camera parameters based on estimated calibration board orientation from different cameras, we found that this is often not sufficient to obtain accurate camera calibrations, especially when there are few boards detected. To resolve this issue, we implemented procedures that optimize the camera calibration parameters to minimize the reprojection error of the calibration board keypoints, referred to as bundle adjustment in the camera registration literature [24]. We implemented bundle adjustment with standard (least-squares) as well as robust losses (Huber and soft L1). Furthermore, we developed an iterative procedure we term “iterative bundle adjustment”, which performs bundle adjustment in multiple stages, using only a random subsample of detected keypoints points in each stage (see Methods for a detailed description). This procedure automatically tunes the outlier thresholds and minimizes the impact of erroneous keypoint detections and bad camera initialization. Each of these bundle adjustment procedures improves the reprojection error from the initial estimate (Figure 3B). Iterative bundle adjustment produced marginally better results, but with no parameter tuning, so we use this as the default in Anipose.

### 2.2 Accurate reconstruction of physical lengths and angles in 3D

An important test of any calibration method is whether it can accurately reconstruct an object with known dimensions. We evaluated the Anipose calibration and triangulation toolkit by asking whether it could estimate the lengths and angles of a precisely manufactured ChArUco board [25].

We first compared the accuracy of tracking the 9 corners of the ChArUco board (Figure 4A) with three methods: manual annotation, neural network detections, and OpenCV detections (example detections in Figure 4B). Although manual annotations are typically assumed to be the ground truth in tracking animal kinematics, we started by assessing the reliability of manual annotations relative to high-precision, sub-pixel resolution keypoint detection based on the geometry of the ChArUco board with OpenCV [23, 25]. Relative to the OpenCV points, the manual keypoint annotations had a mean error of (0.52, −0.75) pixels and standard deviation of (2.57, 2.39) pixels, in the (x, y) directions, respectively (Figure 4C). These observations provide a useful baseline of manual annotation accuracy.

**Figure 4:**
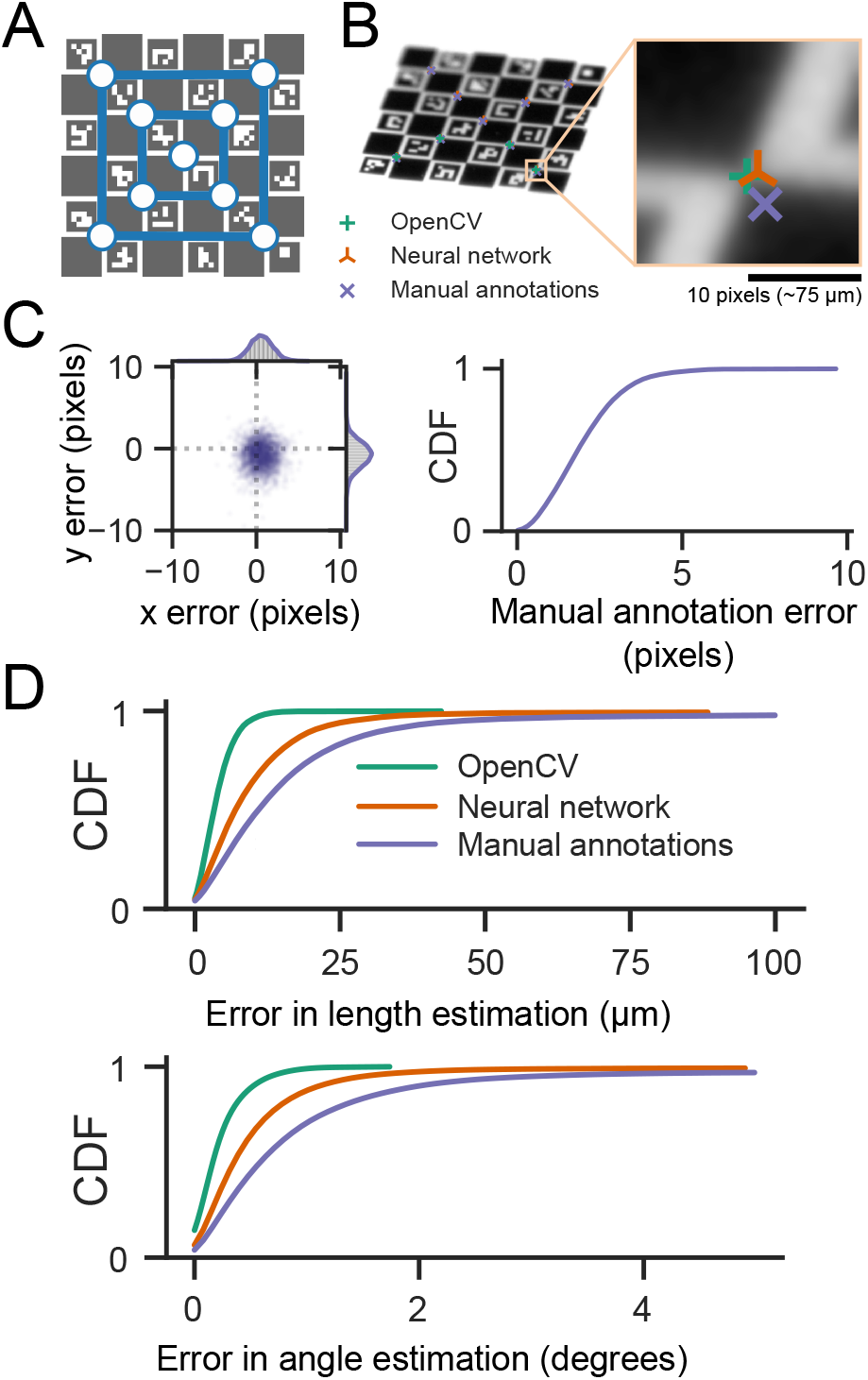
Anipose reliably estimates edge lengths and angles of a precision manufactured ChArUco calibration board. (A) We identified 9 corners as keypoints on the ChArUco board in 200 frames from each of 6 cameras. (B) For comparison, we used manual annotation of the same ChArUco board dataset to train a neural network. We then compared tracking errors of the manual annotations, the neural network, and OpenCV. (C) Error in manually annotated keypoints relative to the sub-pixel precision of OpenCV detections. Manually annotated keypoints had a mean error of (0.52, -0.75) pixels and standard deviation of (2.57, 2.39) pixels. (D) Lengths between all possible pairs of keypoints were computed and compared to the physical lengths. Similarly, all possible angles between triplets of keypoints were computed and compared to known physical angles. OpenCV keypoints provided the most reliable estimates, followed by neural network predictions, then manual annotations. Note that OpenCV generally detected only a small fraction of the keypoints detected by the neural network or through manual annotation (19.3% of keypoints detected by OpenCV, compared to 78.1% by the neural network and 75% by manual annotations).

We evaluated the accuracy of reconstructing ChArUco board lengths and angles as estimated by three methods: manual keypoint annotations, OpenCV keypoint detections, and neural network keypoint detections (see Methods for detailed descriptions). As our ground-truth dataset, we chose the known physical lengths and angles between all pairs of 9 corners on the ChArUco board. The ChArUco board was manufactured with precise tolerance (*<* 2 µm), which allowed us to evaluate the accuracy of lengths and angles from manual keypoint annotations and OpenCV keypoint detections, which are commonly taken to be the ground truth. As expected, OpenCV detections had the lowest error in length and angle, as they leveraged prior knowledge of the ChArUco board geometry to make high-precision corner estimates (Figure 4D). Surprisingly, neural network (trained with DeepLabCut) predictions had a lower error than manual annotations, despite the network itself being trained on manual annotations. More than 90% of poses estimated by Anipose had an error of less than 20 *µ*m in length and 1 degree in angle, relative to the true dimensions of the ChArUco board (Figure 4D). These results demonstrate the efficacy of camera calibration with Anipose and serve as useful bounds of expected performance.

### 2.3 Animal tracking in 3D

We evaluated the triangulation of markerless tracking on three different animal datasets (Figure 5). For each dataset, we computed the error of estimated joint positions and angles on labeled animals withheld from the training data. The error in estimated joint angles was *<*16° in over 90% of frames, and *<*10° in over 75% of frames. Furthermore, the error in the estimated joint position was *<*18 pixels (approximately 1.6mm, 0.14mm, 86mm for mouse, fly, and human datasets respectively) in over 90% of frames and *<*12 pixels (approximately 1mm, 0.09mm, 57mm for mouse, fly, and human datasets respectively) in over 75% of frames. Importantly, the position error in units of camera pixels is roughly comparable across these three datasets, spanning more than 3 orders-of-magnitude in spatial scale. Therefore, we believe these errors are representative of what can currently be expected for accuracy of 3D marker-less tracking.

**Figure 5:**
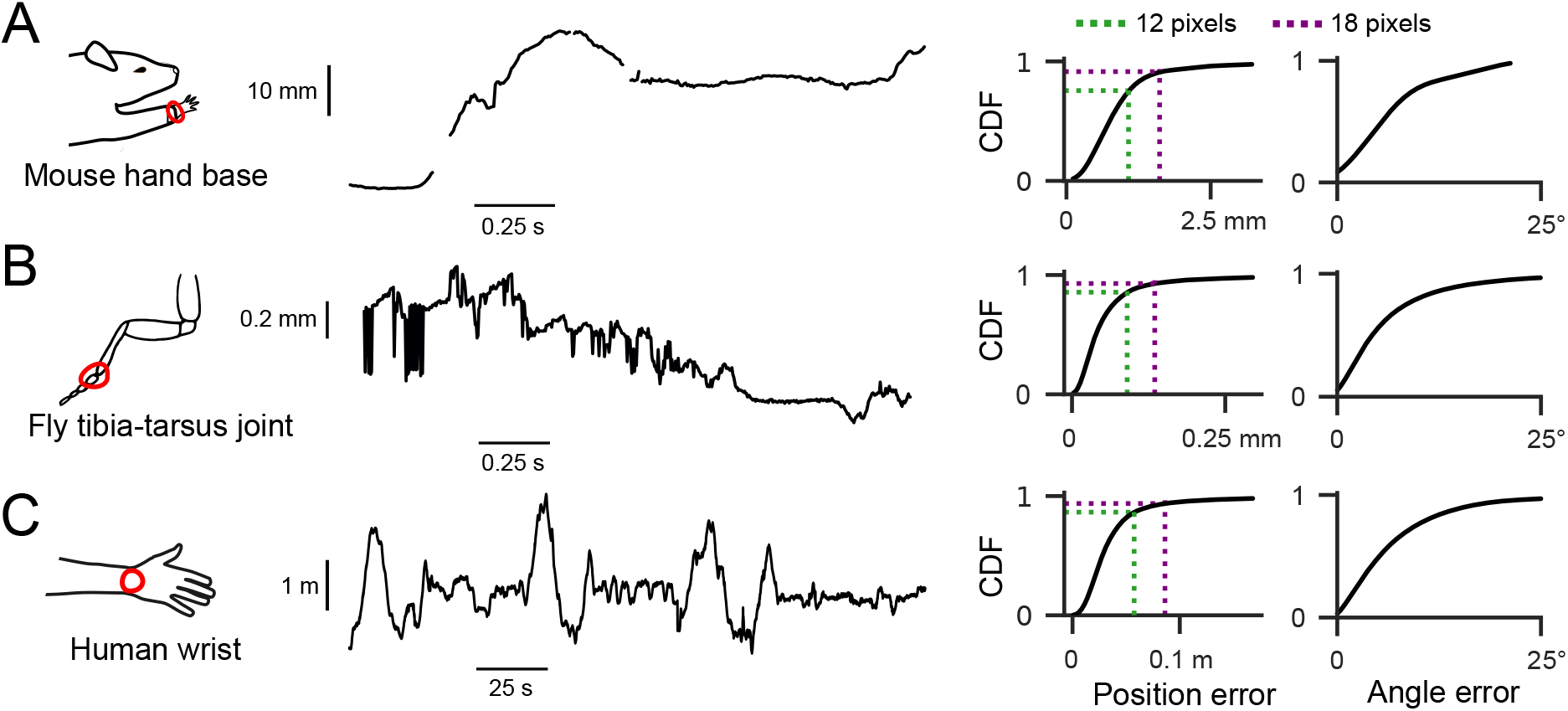
Anipose can consistently estimate positions and angles of joints across three different datasets. In each dataset, the position error is below 12 pixels for 75% of the frames, and below 18 pixels for 90% of the frames. At this stage, prior to filtering, outlier and missing keypoint detections are apparent. (A) Shown at left is an example trace of the tracked 3D position of the base of the mouse hand, projected onto the direction of the reach. On the right, we quantified the distribution of errors when estimating all joint positions and angles, relative to manual annotations. For the mouse dataset, 1 pixel corresponds to approximately 0.09 mm. (B) Same layout as A, but for 3D position of the fly hind-leg tibia-tarsus joint, projected onto the longitudinal axis of the fruit fly. For the fly dataset, 1 pixel ≈.0075 mm. (C) Same layout as A, but for tracked 3D position of a human wrist, projected onto an arbitrary axis. Note that the human (and their wrist) is moving throughout the room. For the human dataset, 1 pixel ≈4.8 mm.

Although triangulation usually resulted in accurate estimates of joint positions and angles, there were still some frames where it failed due to missing keypoint detections (as in Figure 5A). In other cases, incorrect keypoint detections led to erroneous 3D joint position estimates (as in Figure 5B). Even though these issues occurred in a small minority of frames, tracking errors are especially problematic for analyzing movement trajectories. For instance, missing estimates complicate the estimation of derivatives, whereas erroneous estimates bias the distribution of summary statistics. To minimize these issues, we leveraged complementary temporal and spatial information within each dataset to refine tracking performance in 3D.

### 2.4 Addition of filters to improve tracking accuracy

Naturally behaving animals present unique challenges for 3D pose estimation. Animals can contort their bodies into many different configurations, which means that each behavioral session may include unique poses that have not been previously encountered, even across multiple animals. Our approach to tackling these challenges is to leverage prior knowledge that animal movements are usually smooth and continuous, and that rigid limbs do not change in length over short timescales. In particular, we developed and implemented a set of 2D and 3D filters that refine keypoints, remove errors in keypoint detections, and constrain the set of reconstructed kinematic trajectories. We demonstrate that both sets of filters work together to significantly improve pose estimation. Here we focus on detailed quantification of these filters in tracking flies and humans, where our datasets included keypoints at every limb joint tracked with at least 4 camera views.

#### 2.4.1 Refining keypoints in 2D

We implemented three distinct algorithms to remove or correct errors in 2D keypoint detection: a median filter, a Viterbi filter, and an autoencoder filter. The median and Viterbi filters operate on each tracked joint across frames, and the autoencoder filter refines keypoints using learned correlates among all joints. The median filter removes any point that deviates from a median filtered trajectory of user-specified length, then interpolates the missing data. The Viterbi filter finds the most likely path of keypoint detections for each joint across frames from a set of top (e.g., 20) detections per frame, given the expected standard deviation of joint movement in pixels as a prior. Finally, the autoencoder filter corrects the estimated score of each joint based on the scores of the other joints, with no parameters set by the user. Where errors in tracking cannot be corrected by filtering, the keypoint is removed altogether, since the missing joint can be inferred from other camera views, but an erroneous keypoint can produce large discrepancies in triangulation. We document the parameters we used to produce results across the paper in Table S1. Anipose users are encouraged to evaluate the effect these filtering parameters may have on their analyses. Depending on the particulars of the experimental setup, including the spatial and temporal resolution of the videos, the parameters may need to be adjusted.

The addition of each filtering step noticeably improved the tracking of fly leg joints (Figure 6A). The median and Viterbi filters both reduced spurious jumps in keypoint position, which may occur if the neural network detects a similar keypoint on a different limb or at another location in the frame. The Viterbi filter is able to remove small erroneous jumps in detected keypoint trajectories while also preserving high frequency dynamics, whereas the median filter may mistakenly identify fast movements as an error and remove them. The autoencoder filter removed detections for keypoints which were typically not visible from a given view, which improved 3D position estimates after triangulation (Figure S3).

**Figure 6:**
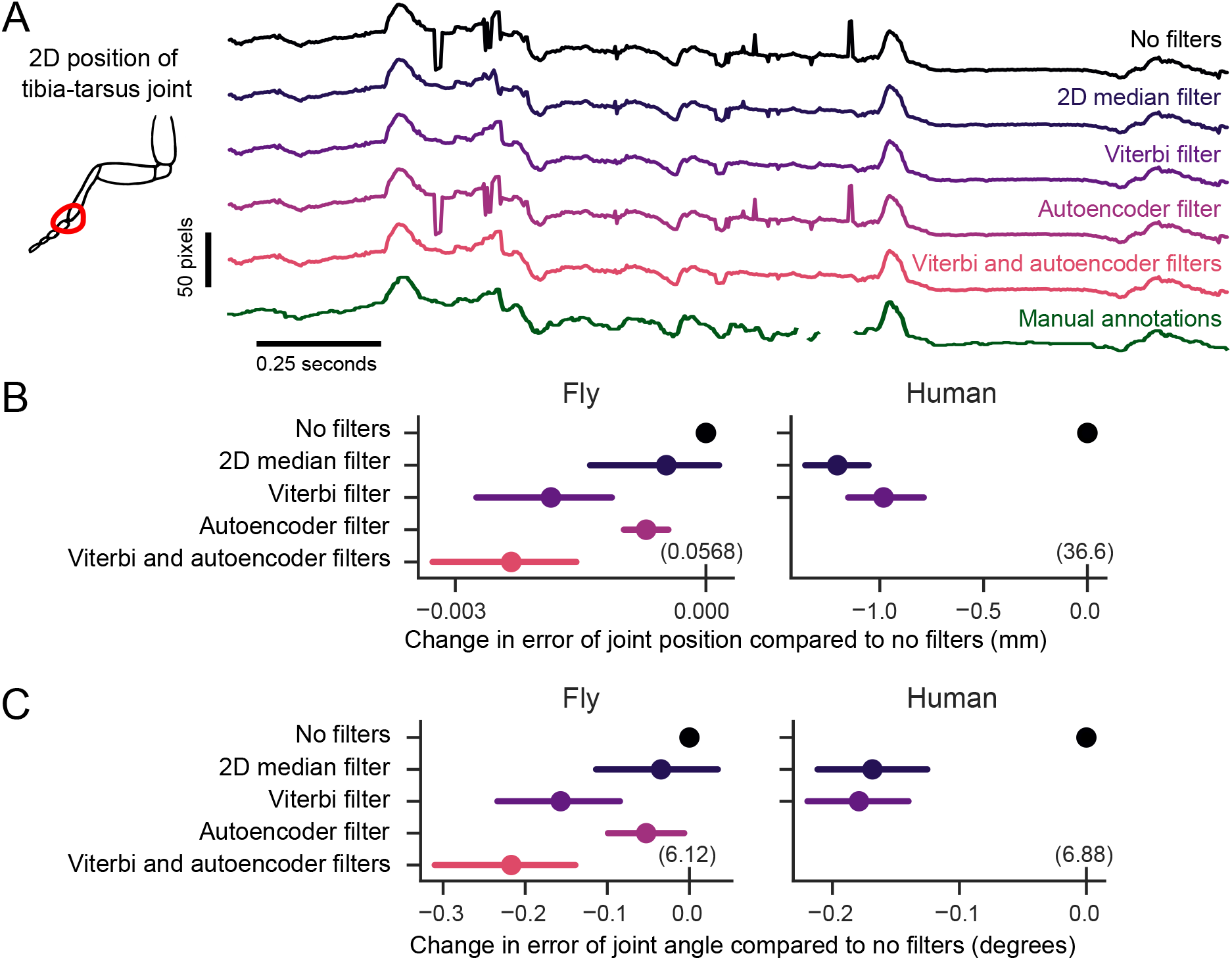
2D filters improve accuracy of 2D pose estimation by taking advantage of the temporal structure of animal behavior. (A) An example trace of the x-coordinate of the 2D position of a fly’s tibia-tarsus joint before and after each step in filtering. Filtering reduces spurious jumps while preserving correct keypoint detections. (B) Comparison of error in joint position before and after filtering. The mean difference in error for the same tracked points is plotted, along with the 95% confidence interval. Viterbi and autoencoder filters significantly improved the estimation of joint position in flies (*p <* 0.001, paired t-test). The Viterbi filter significantly improved estimation of joint position in humans (*p <* 0.001, paired t-test). For the fly dataset, 1 pixel ≈.0075 mm. For the human dataset, 1 pixel ≈4.8 mm. The absolute error values are indicated in parentheses above the 0 tick mark for each dataset. (C) Comparison of angle estimates before and after filtering. The mean difference is plotted as in B. Viterbi and autoencoder filters significantly improved the estimation of angles in flies and humans (*p <* 0.001, paired t-test). The results in (B) and (C) are evaluated on a validation dataset withheld from the training (1200 frames for the fly, 8608 frames for the humans).

For each of the 2D filters, we quantified the performance improvement of estimating the joint position and angle on manually annotated validation datasets. The 2D median filter significantly reduced error in joint position and angle estimation on the human dataset (*t* = −14.8, *p <* 0.001 for position, *t* = −7.7, *p <* 0.001, paired t-test) but not on the fly dataset (*t* = −1.2, *p* = 0.2 for position, *t* = −0.98, *p* = 0.3, paired t-test). The Viterbi filter reduced error on both fly and human datasets (*t* = −4.4 and *t* = −4.1 for fly position and angle, *t* = −10.9 and *t* = −8.7 for human position, with *p <* 0.001 for all, paired t-test). The autoencoder filter also reduced error in joint positions and angles on the fly dataset (*t* = −5.4, *p <* 0.001 for positions, *t* = −2.16, *p* = 0.03 for angles, paired t-test). We did not apply the autoencoder filter to human tracking, since all occluded points are annotated in the training dataset. In the fly dataset, applying the autoencoder filter after the Viterbi filter further improved the joint position and angle estimates above the autoencoder (*t* = −3.97, *p <* 0.001 for positions, *t* = −3.44, *p <* 0.001 for angles, paired t-test). In summary, we found the addition of these three filters improved the ability of Anipose to accurately estimate joint positions and angles.

#### 2.4.2 Refining poses and trajectories in 3D

To further refine joint position and angle estimates in 3D, we developed a novel triangulation optimization that takes advantage of the spatiotemporal structure of animal pose and behavior. Specifically, our optimization produces pose estimates that are smooth in time using temporal regularization, and limbs demarcated by adjacent keypoints that are constant in length with spatial regularization. The length for each limb is automatically estimated in the optimization. The relative strengths of the temporal and spatial regularization terms may be balanced and tuned independently. As with the 2D filters, we empirically determined default strengths that worked across multiple datasets. A complete description of each filter, along with all the parameters, is detailed in the Methods. For illustration, we compared the performance of these filters (Figure 7A) to other commonly used methods from the literature (Random sample consensus, or RANSAC, triangulation and 3D median filter) on the walking fly dataset. We applied the 3D filters on kinematic trajectories partially corrected with 2D filtering (Viterbi then autoencoder filters for the fly dataset, and Viterbi filter only for the human dataset), to evaluate how much the 3D filters improved the accuracy. Spatiotemporal regularization substantially improved pose estimation. The temporal regularization noticeably reduced jitter in the trajectory (Figure 7A), while the spatial regularization stabilized the estimate of limb length (Figure 7B). These improvements are also obvious in example videos of reconstructed pose before and after filtering (Video 2).

**Figure 7:**
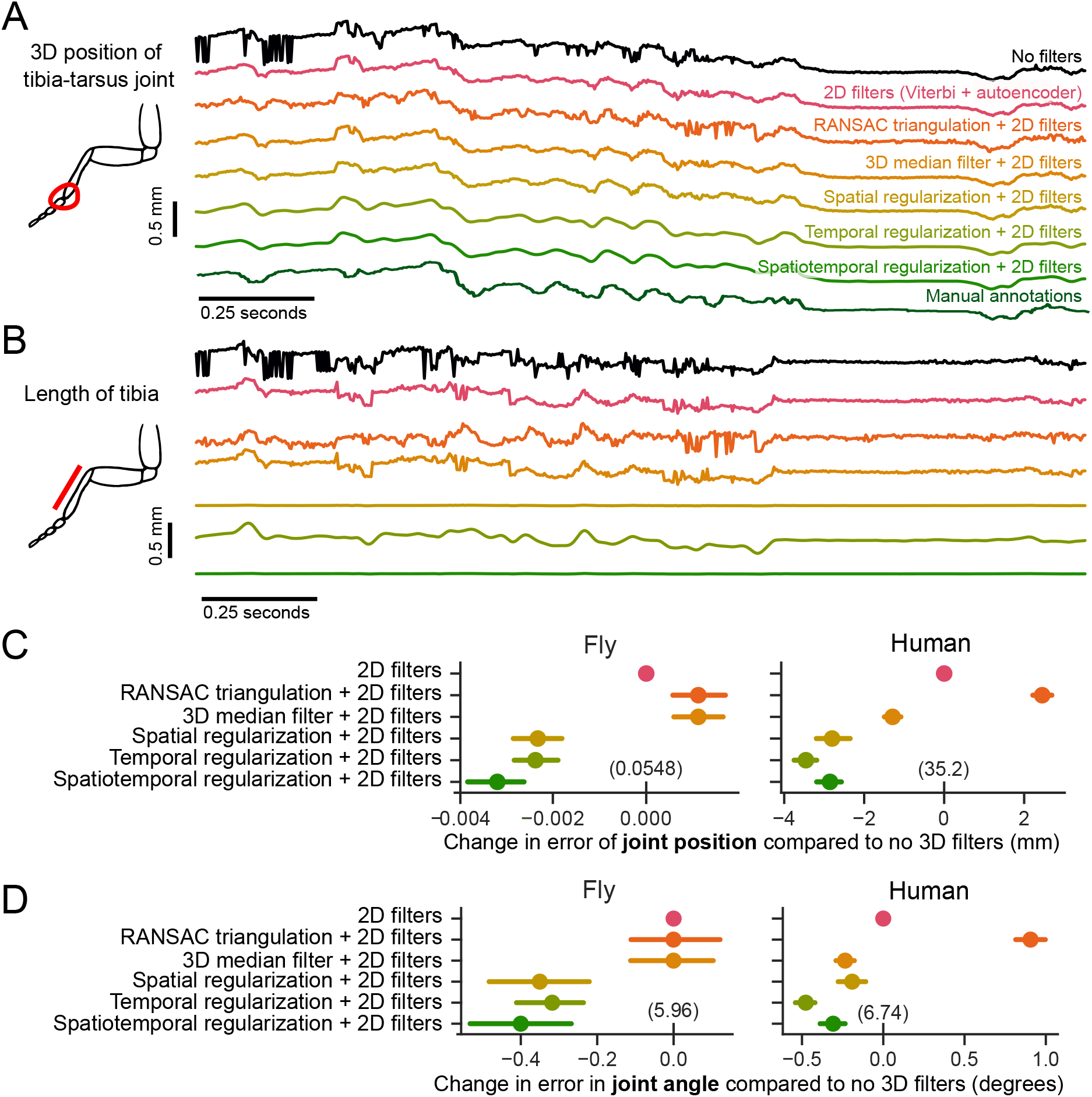
Spatiotemporal filters further improve 3D pose estimation. (A) An example trace of the tracked 3D position of the fly tibia-tarsus joint, before and after filtering. To plot a single illustrative position value, the 3D x-y-z coordinate is projected onto the longitudinal axis of the fly. Also included are comparisons with standard 3D filtering algorithms RANSAC and a 3D median filter, along with manual annotations. Filtering leads to reduction of sudden jumps and keypoint jitters, even compared to 2D filters alone. (B) Estimation of tibia length over time, before and after filtering. Adding spatial regularization leads to a more stable estimate of the tibia length across frames. (C) Comparison of error in joint position before and after filtering. The mean difference in error for the same tracked points is plotted, along with the 95% confidence interval. The absolute error values are indicated in parentheses above the 0 tick mark for each dataset. The 2D filters are the Viterbi filter followed by the autoencoder for the fly dataset and Viterbi filter alone for the human dataset. Spatiotemporal regularization improves the estimation of joint position significantly above 2D filters in both datasets (p < 0.001, paired t-test). The 3D median filter improves pose estimation on the human dataset (p < 0.001, paired t-test) but not on the fly dataset. RANSAC triangulation does not improve pose estimation for either dataset. For the fly dataset, 1 pixel corresponds to 0.0075 mm. For the human dataset, 1 pixel corresponds to 4.8 mm. (D) Comparison of angle estimates before and after filtering. The mean difference and confidence intervals are plotted as in C. Spatial and temporal regularization improve angle estimation above 2D filters on both datasets (p < 0.001, paired t-test). The 3D median filter improves angle estimation on the human dataset (p < 0.001, paired t-test) but not on the fly dataset (p > 0.8, paired t-test). RANSAC triangulation does not improve angle estimation for either dataset.

For each of the 3D filters, we quantified the improvement in position and angle error relative to tracking with 2D filters alone (Figure 7C and D). We found that RANSAC triangulation did not improve position and angle error. The 3D median filter significantly reduced position and angle errors relative to only 2D filters for the human dataset (*t* = −11.8 for position, *t* = −7.3 for angle, *p <* 0.001 for both, paired t-test), but not for the fly dataset. Spatial and temporal regularization applied together provided the largest reduction in tracking error (*t* = −18.7 and *t* = −6.1 for human positions and angles, *t* = −10.8 and *t* = 5.8 for fly positions and angles, *p <* 0.001 for all, paired t-test). Overall, we find that the 3D filters implemented in Anipose significantly improve pose estimation.

#### 2.4.3 Improving estimation of derivatives

In addition to tracking body pose, it is often valuable to track the speed of body movements. We compared the temporal derivative of 3D joint positions estimated with Anipose to the derivative computed from manual annotations (Figure 8) and found both qualitative and quantitative improvements to estimation of body movement speed.

**Figure 8:**
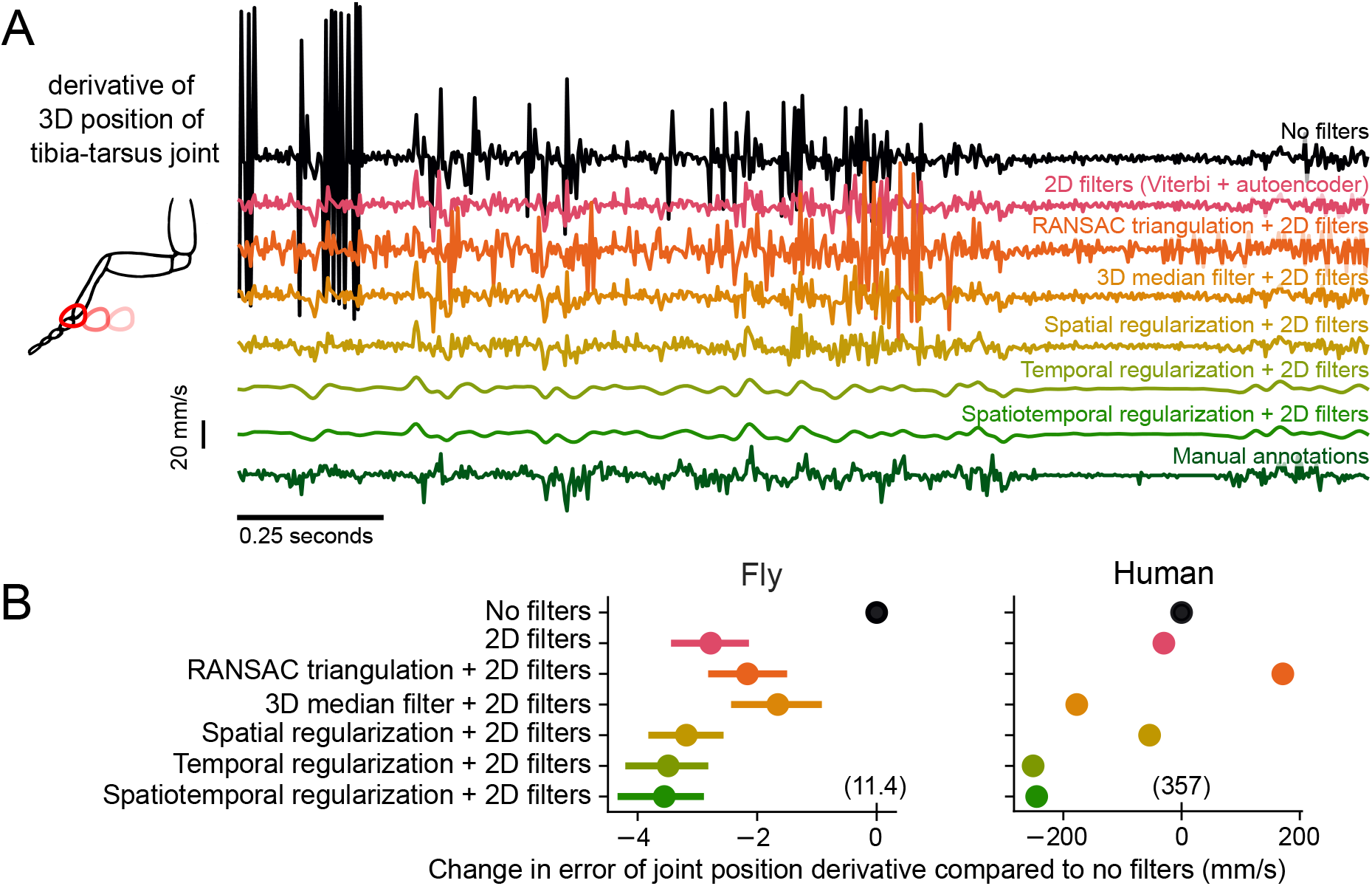
Spatiotemporal filters improve 3D derivative estimation. (A) An example trace of the derivative of the 3D position of the fly tibia-tarsus joint, before and after filtering. To plot a single illustrative derivative value, the 3D x-y-z joint coordinates is projected onto the longitudinal axis of the fly. Spatiotemporal regularization produces smooth derivative estimates, which are closer to the manual annotations compared to other filtering approaches. (B) Comparison of error in joint position derivative before and after filtering. The mean difference in error for the same tracked points is plotted, along with the 95% confidence interval. The absolute error values are indicated in parentheses above the 0 tick mark for each dataset. The 2D filters are the Viterbi filter followed by the autoencoder for the fly dataset and Viterbi filter alone for the human dataset. For the human dataset, due to the large number of labeled points, the confidence intervals are smaller than the size of the points. Adding filters significantly improves the estimate of the derivative.

Filtered trajectories produced smoother derivatives, due to the fact that tracking errors are corrected through 2D and 3D filtering, and the temporal regularization explicitly penalizes deviations from smoothness (Figure 8A). It is challenging to evaluate the accuracy of Anipose derivative estimates because computing finite difference derivatives of manual annotations amplifies known errors in these annotations. Given that manual annotations deviate from the ground truth tracking with a standard deviation of at most 3.5 pixels in distance (Figure 4C), we expect computing the finite difference derivative of such annotations to produce derivatives with error of 4.95 pixels (about 0.037 mm corresponding to 11.1 mm/s over one frame in the fly dataset). Therefore, the manual annotations (dark green trace in Figure 8A) do not represent the true derivative, but rather a noisy approximation of the true derivative. The temporally regularized trajectory resembles this estimate of the derivative but is more smooth because of temporal regularization. The strength of this regularization, and the subsequent smoothness of the tracked keypoints, is a parameter that users may fine-tune (see [26] for a systematic way to tune this parameter). We suggest some default values and provide guidance on choosing parameters in the Discussion.

We found that the 2D filters (Viterbi and autoencoder in fly, only Viterbi in human) improved the error in derivative by 2.78 mm/s for the fly dataset (*t* = −9.4, *p <* 0.001, paired t-test) and by 30.0 mm/s on the human dataset (*t* = −28.0, *p <* 0.001, paired t-test) relative to no filters. The 3D median filter improved the error in derivative by 1.65 mm/s in the fly dataset (*t* = −4.8, *p <* 0.001, paired t-test) and by 177.3 mm/s in the human dataset (*t* = −324, *p* ≪ 0.001, paired t-test) RANSAC improved error in the derivative estimate by 2.16 mm/s in the fly dataset (*t* = −7.07, *p <* 0.001, paired t-test) but did not improve the error in the human dataset. The spatiotemporal regularization improved the error in derivative by an additional 0.67 mm/s for the fly dataset (*t* = −4.10, *p <* 0.001, paired t-test) and by 217.7 mm/s on the human dataset (*t* = −213, *p* ≪ 0.001, paired t-test) relative to the 2D filters. Overall, we found that the filters implemented in Anipose significantly improved the estimation of body movement in the fly and human datasets.

### 2.5 Structured processing of videos

Animal behavior experiments are often high-throughput, meaning that large numbers of videos are recorded over many repeated sessions with different experimental conditions. To make the process of 3D tracking scalable to large datasets, we designed a specific file structure (Figure S7) to organize and process behavior videos, configuration files, and calibration data. This file structure also facilitates scalable analysis of body kinematics across individual animals and experimental conditions. For example, the command `anipose analyze` detects keypoints for each video in the project folder, and `anipose calibrate` obtains calibration parameters for all the cameras in all calibration folders. Each command operates on all videos in the project, circumventing the need to process each video individually. In addition, this design allows the user to easily reanalyze the same dataset using different filtering parameters or with different 2D tracking libraries (e.g., to compare DeepLabCut and SLEAP). For the users that prefer to set up their own pipelines, we also package the calibration, triangulation, and filtering functions in a separate library called aniposelib.

### 2.6 Visualization of tracking

The large number of videos and keypoints tracked in many behavior experiments make it challenging to visualize the resulting data. In addition, the large files created with high-speed video often make it impractical to store and visualize an entire dataset on a laptop. To facilitate evaluation and interpretation of data tracked with Anipose, we developed a web-based visualization tool (Figure 9). The tool shows, for a given trial, each camera view, 3D tracking, and 2D projections of the tracked keypoints. The user can speed up and slow down the speed at which the videos play and rotate the tracked keypoints in 3D. By taking advantage of the standardized file structure, the interface provides a dropdown menu to navigate between trials and sessions. The interface also allows the user to annotate the behaviors in each video, which is particularly useful for isolating specific behaviors for further analysis. As this tool is web-based, it may be run on a server, allowing users to preview videos and inspect tracking from any computer. Furthermore, if the server is public, users may easily share links to particular trials with collaborators to point out specific behaviors (example here).

**Figure 9:**
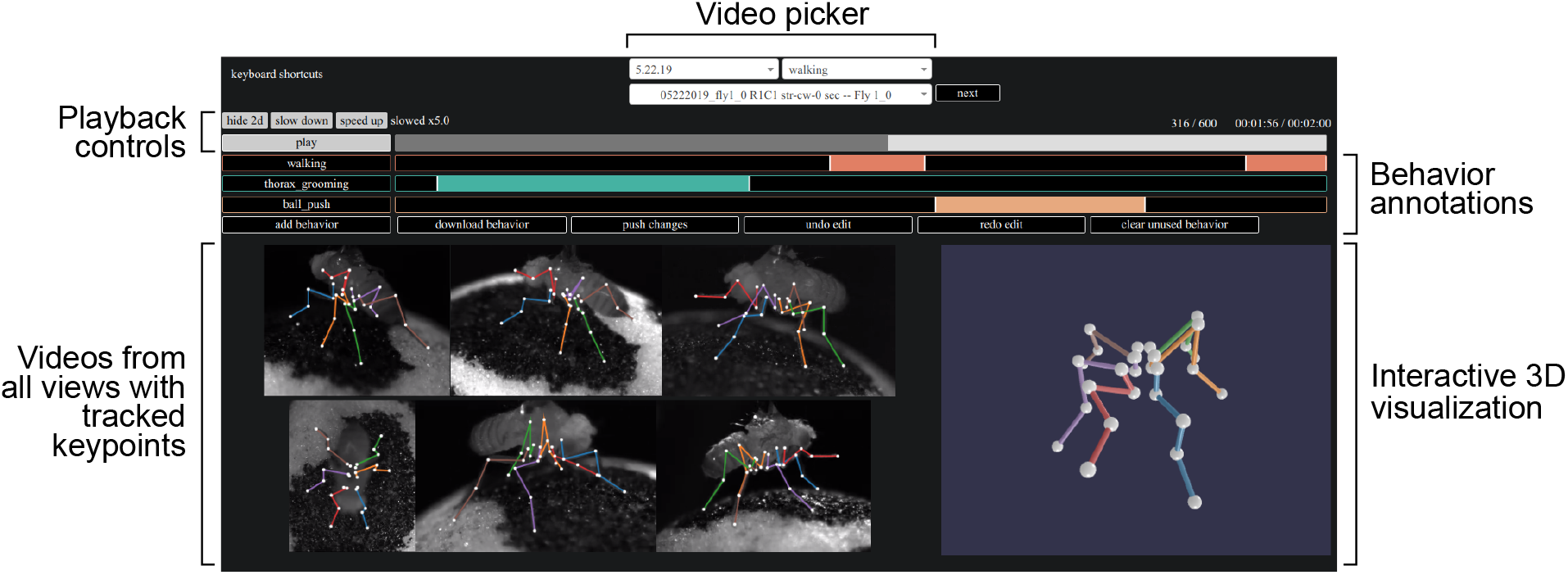
A web tool for visualizing 3D kinematics tracked with Anipose. The videos from all views are displayed synchronously, with overlaid projections of 3D keypoints from Anipose. To the right of the videos, a dynamic 3D visualization allows the user to interact with the 3D keypoints by rotating or zooming in. Above the videos, the user can alter the playback speed or jump to different time points in the video. The user can also annotate the behavior of the animal for further analysis. Menus at the top allow the user to select specific recording dates, experimental trials, or filter trials by a specific behavior.

### 2.7 3D tracking with Anipose provides new insights into motor control of *Drosophila* walking

We first used 3D tracking with Anipose to analyze the leg joint kinematics of fruit flies walking on a spherical treadmill. Although fly walking has been studied in great detail from a 2D perspective [27, 28, 29], 3D joint kinematics of walking flies have not previously been analyzed. Thus, it was not clear how fly leg joints move during walking. Specifically, we sought to understand the relative contributions of leg joint flexion and rotation.

Some limb joints are not restricted to movement in a single plane, but can also rotate around the long axis of a limb segment. Whereas the importance of rotation angles has long been recognized for human gait analysis [30], rotation angles have been comparatively understudied in other animals. This gap exists largely because estimating rotation angles requires precise tracking of joint kinematics in 3D.

The fly leg consists of five segments, whose movements are defined by 8 angles (1 abduction, 3 rotation, 4 flexion). We observed significant rotations between the coxa and femur segments during walking. Figure 10A shows trajectories of coxa rotation, femur rotation, and femur-tibia flexion angles for one walking bout.

**Figure 10:**
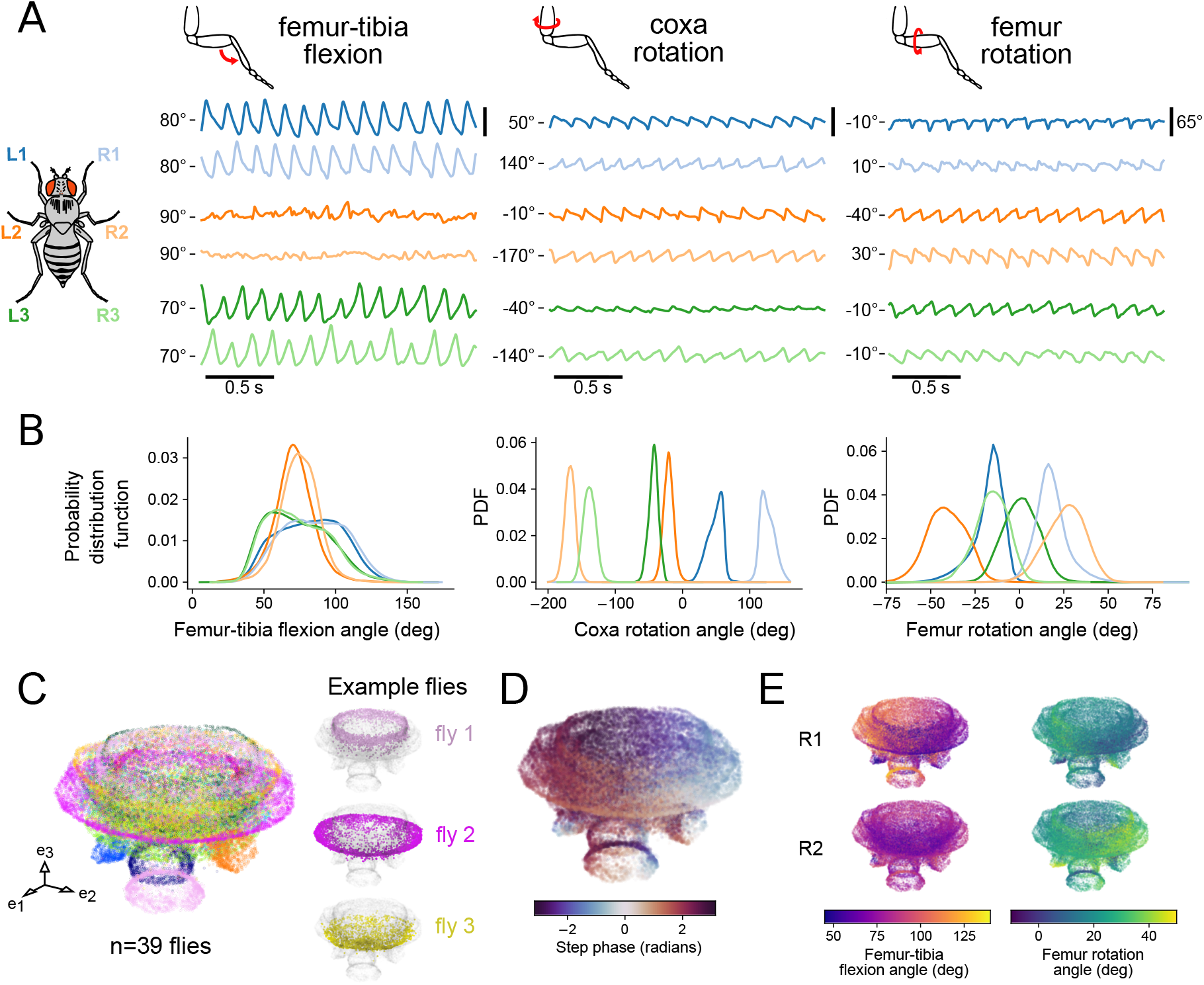
3D tracking of fly walking reveals difference in rotation and flexion angles across legs. (A) Representative traces of coxa rotation, femur rotation, and femur-tibia flexion angles from tethered-walking flies. The median angle value is indicated for each angle as a reference point. (B) Probability distribution functions of coxa rotation, femur rotation, and femur-tibia flexion angles from 39 flies (1480 total seconds of walking). Only walking bouts are included. The distribution of femur-tibia flexion angles is broader for the front and rear legs, whereas the distribution of femur rotation angles is broader for the middle legs. (C) UMAP embedding of coxa rotation, femur rotation, femur-tibia flexion angles across all legs, and their derivatives. Axis units are arbitrary. Although each fly has a characteristic gait, there is a continuum across all flies. (D) UMAP embedding as in C, colored by the phase of the step cycle, revealing the match between the circular structure of the embedding and the step phase. (E) UMAP embedding as in C, colored by front-right leg femur-tibia flexion and femur rotation, and middle right leg femur-tibia flexion and femur rotation. Across multiple flies, the dynamics of the middle legs are dominated by femur rotation, whereas the dynamics of the front legs are dominated by femur-tibia flexion.

Interestingly, the magnitude of joint rotation varied across different legs. Although the femur-tibia flexion angle has a high range of motion in the front and back legs, the femur-tibia flexion angle has a comparatively smaller range of motion in the middle legs (Figure 10B). In contrast, the middle legs are primarily driven by coxa and femur rotation. Furthermore, the coxa joints of contralateral legs rotate in opposing directions. These results suggest that the circuitry that coordinates walking (e.g., the central pattern generator) cannot be the same for all six legs. Rather, walking circuits must control different motor neurons and muscles to generate unique joint kinematics for each leg.

In addition to comparing joint angle distributions across legs, we analyzed trajectories of 3D leg kinematics across flies. We used the UMAP nonlinear embedding method [31] to embed coxa rotation, femur rotation, and femur-tibia flexion angles and their derivatives of all legs (Figure 10C). The three-dimensional embedding of joint kinematics formed a mushroom-shaped manifold. Individual flies reside at specific regions of the manifold, but for all flies, step phase is distributed along the circumference of the cap (Figure 10D). These results are consistent with the existence of a continuum of walking gaits across flies [27], but also suggest that different flies have slightly distinct walking kinematics. This analysis also demonstrates how 3D tracking can be used to dissect the contributions of specific joints to complex motor behaviors. Visualizing a manifold of 3D joint kinematics provides a means to understand how joint kinematics vary within the high-dimensional space of a motor control task (Figure 10E, Figure S8B).

### 2.8 Analysis of 3D mouse reaching and human walking kinematics

To illustrate the value of 3D tracking with Anipose for studying other animal species, we analyzed data from reaching mice and walking humans. Joint positions and angles have long been used to quantify movement in both healthy and impaired animals [32, 33, 34]. However, previous quantification has relied primarily on laborious manual tracking or marker-based tracking with extensive manual corrections. Here we demonstrate analysis of mouse and human behavior using fully automated 3D tracking with the Anipose toolkit.

We first analyzed 3D hand trajectories from mice trained to reach for and grasp a pellet. This task has been extensively used to study neural circuits for sensorimotor control underlying skilled limb movements [35, 36, 37, 38, 39, 40]. Using the Anipose visualization tool, we labeled the reach outcome and start/end frame for each trial. We labeled the trial a “hit” if the mouse successfully grasped the pellet, a “miss” if the mouse missed the pellet holder, and a “bump” if the mouse bumped into the pellet holder or the pellet but failed to grasp the pellet. Each of the four mice in the dataset had multiple instances of each outcome. Figure 11A shows example 3D reaching trajectories, which demonstrate that reaching movements vary significantly from trial to trial (see also Figure S9A). Although reaching is a challenging behavior to track, due to its speed and variability, Anipose was able to accurately reconstruct forelimb reaching trajectories. The trajectory of each movement was variable, but plotting the distance to the pellet holder as a function of time to contact revealed that each reach type has a stereotyped trajectory (Figures 11B and S9B). Interestingly, the hit/bump and miss trajectories diverged around 50 ms prior to pellet contact, suggesting that mice are unable to correct their reaching trajectories in this period.

**Figure 11:**
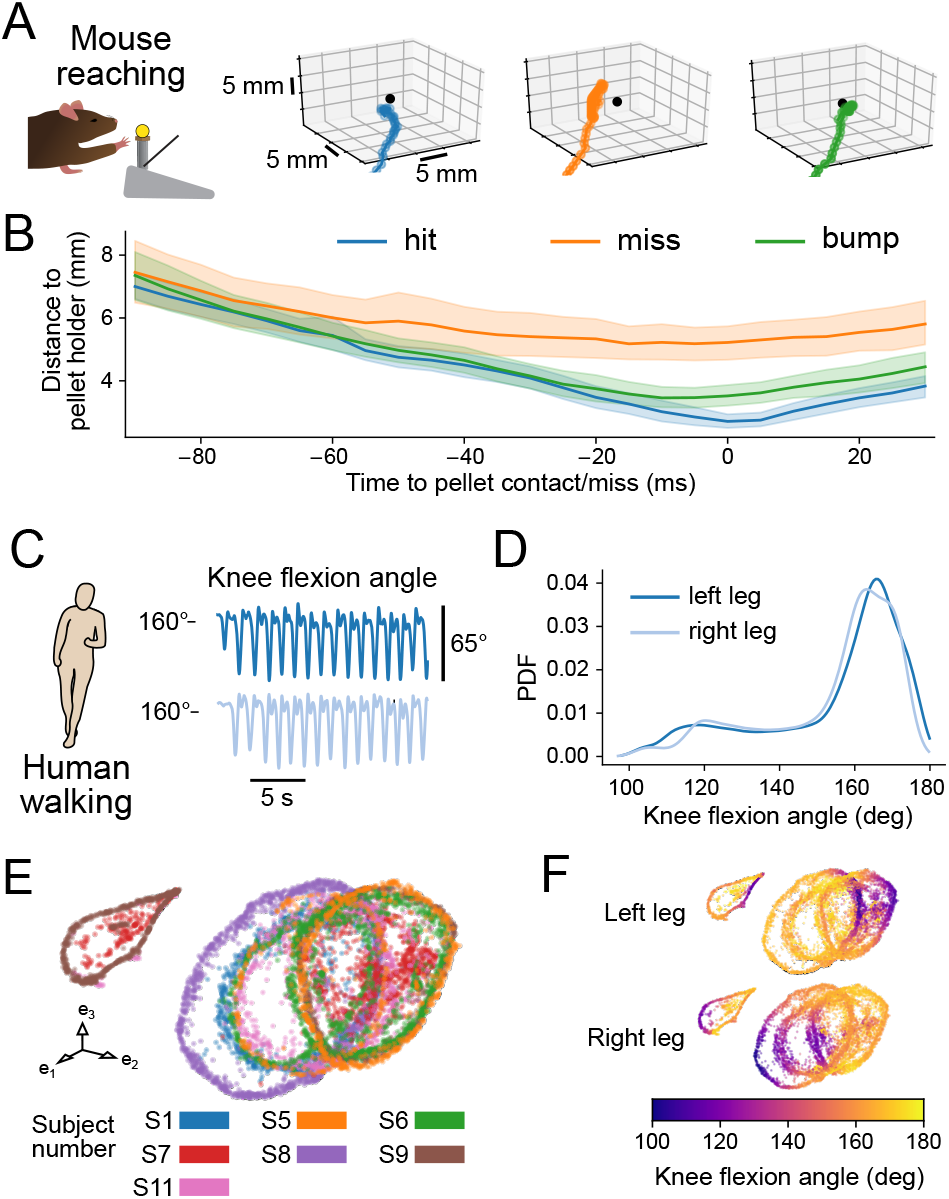
3D tracking with Anipose facilitates analysis of mouse and human kinematics. (A) Example 3D trajectories of a mouse reaching for a food pellet. The pellet is indicated as a black dot. (B) Mean distance to pellet holder as a function of time across all 4 mice (88 hits, 69 bumps, 28 misses). Shaded areas are 95% confidence intervals. When reaches are aligned to the grasp attempt (0 ms), the hand is farther from the pellet holder on miss trials compared to hit or bump trials. Averaging across all mice reveals a clear difference between reach types. (C) Representative trace of knee flexion from a walking human, tracked with Anipose. Data is from the Human 3.6M dataset. The median angle value is indicated at left as a reference point. (D) Probability distribution function of knee flexion angle from 7 humans. Only sessions that include walking are included. (E) UMAP embedding of knee flexion, hip rotation, and hip flexion angles across all legs, and their derivatives. Axis units are arbitrary. Although each human subject has a characteristic gait, there is a continuum across all subjects. (F) UMAP embedding as in E but colored by knee flexion for each leg. Coloring by knee flexion angle reveals the common phase alignment of the circles across subjects.

We next analyzed 3D walking kinematics reconstructed from the human dataset using methods similar to our analysis of fly walking. We extracted knee flexion, hip rotation, and hip flexion angles from 3D joint positions tracked with Anipose (Figures 11C and S10A). The distributions of these joint angles are symmetric across the two legs (Figures 11D and S10B) and match previous characterizations of human gait [34]. To characterize the structure of walking across the subjects, we used the UMAP nonlinear embedding method [31] to embed knee flexion, hip rotation, hip flexion, and their derivatives into a 3D space, as for the fly dataset above. The UMAP embedding reveals a continuous manifold of angle coordination across subjects (Figure 11E). The manifold forms a cylindrical structure with the knee flexion angle mapping circularly along the cylinder (Figure 11F). The two trials that are to the left outside the main cylinder have lower variation of left leg hip rotation (Figure S10C). These examples illustrate the ease and utility of tracking and analyzing human walking behavior with Anipose. In the future, this approach could be used to automatically identify individuals with distinct walking gaits or other motor patterns.

## 3 Discussion

In this paper, we introduce Anipose, a new open-source toolkit to accurately track animal movement in 3D. Anipose is designed to augment DeepLabCut, a toolkit for 2D markerless tracking [11], with calibration, filters, and a visualization tool to facilitate robust 3D tracking and analysis. Current users of DeepLabCut can easily upgrade to 3D tracking with Anipose by adding and calibrating additional cameras to an existing behavioral setup. Although we designed Anipose to leverage 2D tracking with DeepLabCut [11], it can be made compatible with other 2D markerless tracking methods, including SLEAP [12] and DeepPoseKit [13] by modifying a single file. We validated each new optimization module and the full pipeline against ground truth data from four different experimental datasets and three organisms, demonstrating accurate reconstruction of 3D joint positions and angles.

The Anipose tracking pipeline is designed to streamline structured processing of videos recorded in high-throughput experiments. Users do not need to know Python to use the Anipose pipeline. All that is required to get started is editing a small configuration file and running the provided commands from a terminal. We also provide access to individual functions via a separate library, aniposelib. To help new users get started, we developed detailed tutorials for both the Anipose pipeline and aniposelib at anipose.org.

### 3.1 Impact of robust markerless 3D tracking

A key technical advantage of tracking with Anipose is the ability to interpret and analyze movement speed from 3D pose trajectories that are smooth in space and time, due to filtering and interpolation from multiple camera views. The resulting improvements in tracking smoothness make it easier to analyze pose and movement dynamics. Specifically, interpolated data enables the user to obtain better estimates of behavior statistics, such as mean and variance, and to perform dimensionality reduction techniques, such as principal component analysis (PCA). Additionally, temporal regularization reduces noise in the first derivative and thus enables the user to obtain more precise estimates of movement speed (Figures 8 and S5).

This ability to analyze 3D pose trajectories may open up new opportunities for behavioral neuroscience, where key insights have been gained through carefully controlled behavioral paradigms. In particular, experiments are often designed to accommodate the practical limitations of movement tracking, recording neural activity, and perturbing the animal in real time (e.g., [41, 42, 43, 44, 45]). Recent advances in experimental technologies (e.g., high-density extracellular recording probes [46], optical imaging of fluorescent reporters [47, 48], and optogenetics [49]) have made it feasible to precisely record and perturb neural activity from animals behaving freely in three dimensions. Complementing these technologies, a comprehensive toolbox for high-throughput 3D tracking will not only enable deeper analysis of current experiments, but also make it possible to study more natural behaviors.

A robust 3D markerless tracking solution could also greatly expand the accessibility of quantitative movement analysis in humans. Many neurological disorders, including some commonly thought of as cognitive disorders, affect walking gait [50, 51] and upper limb coordination [52, 53]. Many clinicians and basic researchers currently rely on qualitative evaluations or expensive clinical systems to diagnose motor disorders and assess recovery after treatment. While clinical approaches are commercially available [54], they are costly, require proprietary hardware, rely on the addition of markers to the patient, and cannot assess walking gait in natural contexts such as a patient’s home. Anipose could be used as a tool in the diagnosis, assessment, and rehabilitative treatment of movement and neurodegenerative disorders.

### 3.2 New insights into the motor control of *Drosophila* walking

By analyzing 3D joint kinematics of tethered walking *Drosophila*, we found that each leg has a unique set of joint angle distributions. One valuable insight, which was not evident from 2D tracking alone, is that the movement of the middle legs is driven primarily by femur rotation, in contrast to the front and hind legs, which are driven primarily by femur-tibia flexion. We also observed small differences in femur-tibia flexion and femur rotation distributions between front and hind legs (Figure 10B). Thus, the neural circuits that move each leg during walking must be specialized for controlling joints with distinct forces and dynamics within each leg. Previous models of *Drosophila* walking have used an identical control architecture for intra-leg joint coordination for all six legs [55, 56]. Our results provide a framework for constructing more biologically plausible neuromechanical models using distinct architectures for controlling different joints within each leg.

Inter-leg differences in joint kinematics also raise new questions about limb proprioception. Proprioceptors in the fly femoral chordotonal organ (FeCO) encode femur-tibia flexion and movement [57]. Does the role of the FeCO differ for the middle legs, for which the femur-tibia generally does not flex in a rhythmic pattern during walking? Which proprioceptors, if any, are used to sense femur and coxa rotation of the middle legs? Answering these questions will be facilitated by combining Anipose with *in vivo* measurements and perturbations of proprioceptive neural circuits [58].

Rythmic motor behaviors, such as walking, are thought to be controlled by central pattern generators (CPGs): neural circuits that generate intrinsic rhythmic activity [59]. If fly walking is controlled by CPGs, our results suggest that the CPG for each leg must control different muscles. For example, we would predict that a walking CPG for the front legs would connect to motor neurons that control the tibia flexor and extensor muscles in the femur [60]. In contrast, a CPG for the middle legs might connect to motor neurons innervating muscles in the trochanter that control femur rotation. These insights will be useful in guiding ongoing efforts to trace motor control circuits using connectomic reconstruction of the *Drosophila* ventral nerve cord [61] and leg [62].

Femur rotation is also likely to be important for walking in other insect species. Fransevich and Wang tested the passive rotation of the trochanter-femur articulation in 23 insect species and found rotation ranges from 10°to 120°, depending on the species [63]. Our estimate for the physiological range for walking *Drosophila* is about 70°(Figure 10B), which falls within the trochanter-femur articulation range observed in other insects. Thus, it is plausible that articulation of the trochanter-femur joint is sufficient to account for the femur rotation we measured during walking, and that other insects rely on femur rotation during walking as well. As an example, Bender et al. reported different kinematics across legs in walking cockroaches, with larger femur rotation and smaller femur-tibia flexion in the middle legs relative to the hind legs [4]. The application of Anipose to track 3D joint kinematics in other species will enable further comparative studies of the biomechanics and neural control of walking.

### 3.3 Potential for future improvement based on related work

Camera calibration has long been a rich topic in computer vision research. The most commonly used calibration code, based on Zhang’s work [64] and part of OpenCV [23], can calibrate up to 2 cameras using images of checkerboards from multiple angles. Although this method can be used to calibrate 3 or more cameras by calibrating pairs of cameras, in practice, precise calibration requires an additional optimization step called bundle adjustment [24]. Bundle adjustment has been a key part of structure from motion toolkits [65, 66], but the method has received comparatively little attention as a solution to camera calibration for markerless tracking. An exception is DeepFly3D, which supports calibration based on animal keypoints but not based on a calibration board, which hinders its ability to handle setups with arbitrary camera positions [20]. Our key innovation is to provide an open source implementation of sparse bundle adjustment targeted for camera calibration for motion tracking. Our current implementation could eventually benefit from incorporating other methods from the literature. For instance, using a neural network to detect the calibration board may yield more detected keypoints (Figure 4) and lead to more robust calibration under difficult conditions [67]. Currently, Anipose requires a calibration board to initialize camera parameters (even with animal calibration), but it may be possible to initialize camera parameters based on commonly detected points, as is commonly done in the structure from motion literature [65, 66], or perhaps by using a neural network directly [68]. Bundle adjustment itself may be made more robust by incorporating gauge constraints in the optimization function, further reducing the number of parameters [24]. Finally, the calibration process itself may be streamlined if it were made interactive [69].

There has been extensive recent work to improve markerless tracking based on deep learning approaches. One common approach has been to improve the neural network architecture for training. For instance, this approach has been used to induce priors in the neural network based on occlusions [70, 71], multi-view geometry [72, 19, 18, 73], limb lengths [74], or time [75]. We note that this approach is complementary to our work, as the Anipose filters could be used with keypoint detection by any neural network. Another approach is to resolve tracking by using pictorial structures to add priors on limb lengths [76, 77, 20] or motion [78]. The Viterbi filter used in Anipose is analogous to the motion based pictorial structures and could be further extended to handle priors on limb lengths based on insights from these papers. Beyond tracking single animals, toolboxes like SLEAP [79], OpenPose [14], and DeepLabCut [22] have some support for multi-animal pose estimation in 2D. For tracking multiple animals in 3D, a promising approach is to build correspondences based on geometry and appearance [80] across multiple views. As automated, high-throughput tracking of animal behavior grows in scale, new methods for data analysis, visualization, and modeling will also be needed to gain insight into the neural control of dynamic behavior [81, 10, 45, 58].

### 3.4 Limitations and practical recommendations

There are several common scenarios under which Anipose may fail to produce accurate 3D tracking. Below, we enumerate some of the scenarios we have encountered in applying Anipose on different datasets and suggest practical strategies for troubleshooting.

As is the case for any tracking system, the ability of Anipose to track and estimate body pose is fundamentally limited by the quality of the underlying data. High quality videos are well illuminated, contain minimal motion blur, and provide coverage of each keypoint from different views. A common failure mode we encountered was when the neural network misplaced 2D keypoints in some frames. If the errors are uncorrelated across camera views, then the Anipose filters can compensate and still produce accurate tracking in 3D. But in some cases, multiple views have correlated errors or these errors persist in time. These type of errors most commonly arise when the neural network has not been trained on a subset of rare behaviors, so that the animal adopts poses unseen by the trained network. One solution to reducing the frequency of such errors involves systematically identifying outlier frames, manually relabeling them, then retraining the network. Anipose supports this functionality, as do other tracking toolboxes [11, 79, 13, 20].

Poor multi-camera calibration also results in tracking errors. A good calibration should have an average reprojection error of less than 3 pixels, and ideally less than 1 pixel. To obtain a quality calibration, the calibration videos should be recorded so that the board is clearly visible from multiple angles and locations on each camera. If it is not possible to achieve this, we suggest exploring a preliminary calibration module in Anipose that refines an initial calibration based on the detected points on the animal itself. This module was inspired by the animal based calibration in DeepFly3D [20], but our implementation uses the initial calibration from a calibration board as a starting guess, permitting generalization in different setups. It also takes advantage of our iterative calibration procedure to yield robust calibration even with errors in tracking.

An effective experimental setup needs to have an appropriate number of cameras to track all keypoints across possible pose configurations. In particular, each joint must be visible from at least 2 cameras at all times. Thus, for tracking multiple limbs or body parts, we recommend at least 3 equally spaced cameras, so that half of the body is visible from any single camera. We evaluated this quantitatively in the human dataset (Figure S6), where there is a dramatic reduction in error from 2 to 3 cameras.

The mouse reaching dataset is one example where tracking was reasonably accurate without filters, but filters did not further improve tracking accuracy. There are several potential explanations for this result. The reaches are very short (about 40-100 frames or 200-500ms) and the hand is hard to see when it is on the ground, so temporal filters such as the Viterbi filter or temporal regularization lack the information to resolve tracking errors. There are very few keypoints (only 3 per hand) and these can change in distance relative to each other, so the spatial regularization cannot impose strong constraints. With only 2 cameras, the spatiotemporal regularization cannot fully leverage multiple views to remove outliers (Figure S6) and the autoencoder has limited utility. In this situation, using basic linear least-squares triangulation works well enough for analysis (Figure 11A and B). The accuracy of tracking mouse reaching might be improved by labeling more keypoints on each hand, increasing the camera frame rate, and adding more cameras.

As a practical starting point, we recommend users start with no filters to first evaluate the quality of the tracking. If outliers or missing data impede data analysis, then we recommend enabling the default filter parameters in Anipose, which we have found to produce good tracking results across multiple datasets. In some cases, some additional tuning of parameters may be required, especially on datasets with unique constraints or when studying behaviors with unusual dynamics. If any joints are not visible for an extended period of time in certain videos, we recommend disabling the spatiotemporal optimization, as it can hallucinate trajectories, increasing overall error (as in Figure S6). We provide suggestions for tuning parameters in our documentation at anipose.org.

### 3.5 Outlook

We designed Anipose to make markerless 3D tracking simple and broadly accessible for the scientific community. With this goal in mind, we built Anipose on DeepLabCut, a widely used 2D tracking toolkit. As many labs develop machine learning tools for behavior tracking and analysis, we advocate for pooling efforts around common frameworks that emphasize usability [82, 83]. In particular, we suggest that tools be built in a modular way, so that code can be extended and reused in other frameworks. We hope that the Anipose toolkit contributes to these community efforts. We welcome contributions to improve and extend the Anipose toolkit and conversely are ready to contribute the ideas and code from Anipose to other toolkits.

## 4 Methods

### 4.1 Video collection and annotation

#### ChArUco dataset

To evaluate the performance of Anipose compared to physical ground truth, we collected videos of a precision-manufactured ChArUco board [25]. The ChArUco board was manufactured by Applied Image Inc (Rochester, NY) with a tolerance of 2 µm in length and 2° in angle. It is a 2 mm × 2 mm etching of opal and blue chrome, on a 5 mm ×5 mm board. The ChArUco pattern itself has 6 × 6 squares, with 4 bit markers and a dictionary size of 50 markers. With these parameters, the size of each marker is 0.375 mm and the size of each square is 0.5 mm. We filmed the ChArUco board from 6 cameras (Basler acA800-510µm) evenly distributed around the board (Figure 1A), at 30Hz and with a resolution of 832 × 632 pixels, for 2-3 minutes each day over 2 separate days. While filming, we manually rotated the ChArUco board within the field of view of the cameras. These videos were used as calibration videos for both the ChArUco dataset and the fly dataset detailed below.

We chose 9 of the corners as keypoints for manual annotation and detection (Figures 1A and 4A). We extracted and manually annotated 200 frames from each camera from day 1, and an additional 200 cameras per camera from day 2 (1200 frames per day, 2400 frames total). We used the frames for day 1 for training the neural network and the frames from day 2 for evaluation of all methods.

#### Mouse dataset

Reaching data were obtained from four adult C57BL/6 mice (∼8-12 weeks old, two male and two female) trained to reach for a pellet. Procedures performed in this study were conducted according to US National Institutes of Health guidelines for animal research and were approved by the Institutional Animal Care and Use Committee of The Salk Institute for Biological Studies. The reaching task is described in detail elsewhere [35]. Briefly, the training protocol consisted of placing the mouse in a 20 cm tall ×8.5 cm wide ×19.5 cm long clear acrylic box with an opening in the front of the box measuring 0.9 cm wide and 9 cm tall. A 3D-printed, 1.8 cm tall pedestal designed to hold a food pellet (20 mg, 3 mm diameter; Bio-Serv) was placed 1 cm away from the front of the box opening and displaced to one side by 0.5 cm (to encourage mice to use their preferred forelimb), and food pellets were placed on top as the reaching target (Fig. 1B). Mice were food deprived to ∼85% of their original body weight and trained to reach for food pellets for either 20 minutes or until 20 successful reaches (defined as pellet retrieval) were accomplished. Mice were trained in this setup for 14 consecutive days before reaches were captured with 2 cameras (Sentech STC-MBS241U3V with Tamron M112FM16 16mm lens) placed in front and to the side of the mouse (∼85° apart). Videos were acquired at a frame rate of 200 Hz at a resolution of 1024 × 768 pixels.

We chose 6 points on the mouse hands as keypoints (Figure 1B). On each mouse hand, we labeled 3 points: the dorsal wrist, the base of digit 5, and the proximal end of digit 3. In total, we manually labeled 2200 frames (1100 frames per camera) for training the neural network from 2 mice. For test data to evaluate the post estimation performance, we labeled an additional 400 frames (200 frames per camera) taken from videos of 2 mice that were not in the training set.

#### Fly dataset

We next evaluated 3D tracking with Anipose on walking fruit flies. Male and female Berlin wild type *Drosophila melanogaster*, 4 days post-eclosion, were used for all experiments. Flies were reared on standard cornmeal agar food on a 14 hr/10 hr light-dark cycle at 25 °C in 70% relative humidity. The flies’ wings were clipped 24-48 hours prior to the experiment in order to increase walking and prevent visual obstruction of the legs and thorax. For all experiments, a tungsten wire was tethered to the dorsal thorax of a cold-anesthetized fly with UV cured glue. Flies were starved with access to water for 2–15 hours before they were tethered. After 20 minutes of recovery, tethered flies were positioned on a frictionless spherical treadmill [84, 85] (hand-milled foam ball, density: 7.3 mg/mm^3^, diameter: 9.46 mm) suspended on a stream of compressed air (5 L/min). Six cameras (imaging at 300 Hz, Basler acA800-510µm with Computar zoom lens MLM3X-MP) were evenly distributed around the fly, providing full video coverage of all six legs (Figure 1C). Fly behavior was recorded in 2 second trials, capturing a range of behaviors such as walking, turning, grooming, and pushing against the ball. The recording region of each video was cropped slightly so that the fly filled the frame and the camera was able to acquire at 300Hz. For all training and test evaluation data, the interval between trials was 25 seconds. For some of the flies in the larger walking dataset used in Figure 10, the interval between trials was set to 9 seconds.

We selected 30 points on the fly as keypoints (Figure 1C). On each fly leg, we labeled 5 points: the body-coxa, coxa-femur, femur-tibia, and tibia-tarsus joints, as well as the tip of the tarsus. In total, we manually labeled 6632 frames (about 1105 frames per camera) for training the neural network. For test data to evaluate the post estimation performance, we labeled an additional 1200 frames (200 frames per camera) taken from videos of 5 flies that were not in the training set. For analyzing flexion and rotation of angles during walking in Figure 10, we used a larger dataset of videos from 39 flies, all collected with the methods described above.

#### Human dataset

We evaluated 3D tracking with Anipose on the Human 3.6M dataset [86, 87]. Because this dataset has been used extensively for human pose estimation, it provides a useful comparison to existing computer vision methods. It consists of 11 professional actors performing a range of actions, including greeting, posing, sitting, and smoking. The actors were filmed in a 4m × 3m space with 4 video cameras (Basler piA1000) at a resolution of 1000 × 1000 pixels at 50Hz (Figure 1D). To gather ground-truth pose data, the actors were also outfitted with reflective body markers and tracked with a separate motion capture system, using 10 Vicon cameras at 200 Hz. Leveraging these recordings, the authors derived the precise 3D positions of 32 body joints and their 2D projections onto the videos. For camera calibration, we used the camera parameters from the Human 3.6M dataset, converted by Martinez et al. [88].

To compare the performance of Anipose against previous methods, we used a protocol from the literature [72]. The Human 3.6M dataset contains data from 5 subjects as a training dataset (2 female and 3 male), 2 subjects as a validation dataset, and 4 subjects as a testing dataset (2 female and 2 male). We used frames from the training dataset to train the network and evaluated the predictions on the validation dataset. We also removed frames from the training dataset in which the subject did not move relative to the previous frame (*<* 40mm movement of all joints from the previous frame). We evaluated the tracked human dataset on every 64th frame. We used 17 of the 32 provided joints as keypoints (Figure 1D). Iskakov et al. [72] showed that some scenes from the S9 validation actor (parts of the Greeting, SittingDown, and Waiting actions) have ground-truth shifted in global coordinates compared to the actual position [72], so we exclude these scenes from the evaluation set. Furthermore, for subject S11, one of the videos is corrupted (part of the “Directions” action), so we exclude this from the dataset as well. In total, we obtained 636,724 frames (159,181 per camera) for training the neural network, and 8608 frames (2152 per camera) frames for evaluation.

#### Manual annotation of datasets

To produce neural network training data, we annotated the fly dataset using Fiji [89] and the VGG Image Annotator (VIA) [90, 91]. All the images in the fly test set were annotated with VIA. We annotated all the images in the ChArUco dataset and mouse dataset with VIA.

### 4.2 Neural network keypoint detections

Detection of keypoints in each of the datasets was performed with DeepLabCut 2.1.4 [22]. Briefly, to produce training data, we used k-means clustering to pick out unique frames from each of the views, then manually annotated the keypoints in each frame. We trained a single Resnet-50 [92] network for all camera views for the fly, mouse, and ChArUco datasets, starting from a network pretrained on Imagenet. For the human dataset, we started with a Resnet-101 network pretrained on the MPII human pose dataset [93]. During training, we augmented the training dataset with cropping, rotation, brightness, blur, and scaling augmentations using Tensorpack [94]. We then used the Anipose pipeline to run the network on each video. For each keypoint, the network produced a list of predicted positions, each associated with a confidence score (between 0 and 1). We saved the top-n most likely predictions of each joint location for each frame for use in Viterbi filtering of likely keypoints in 2D, as described below.

### 4.3 Filtering of 2D keypoint detections

The raw keypoint detections obtained with DeepLabCut were often noisy or erroneous (Figure 6). Thus, filtering the detections from each camera was necessary before triangulating the points. Anipose contains 3 main algorithms to filter keypoint detections; we elaborate on each algorithm below. Example applications of these filters and results are compared in Figure 6.

#### Median filter

The first algorithm identifies outlier keypoint detections by comparing the raw detected trajectories to median filtered trajectories for each joint. We started by computing a median filter on the detected trajectory for each joint’s x and y positions, which smooths the trajectory estimate. We then compared the offset of each point in the raw trajectory to the median filtered trajectory. If a point deviated by some threshold number of pixels, then we denoted this point as an outlier and removed it from the data. The missing points were then interpolated by fitting a cubic spline to the neighboring points. The median filter is simple and intuitive, but it cannot correct errors spanning multiple frames.

#### Viterbi filter

To correct for errors that persist over multiple frames, we implemented the Viterbi algorithm to obtain a single most consistent path in time from the top-n predicted keypoints in each frame for each joint. To be specific, we expressed this problem as a hidden Markov model for each joint, wherein the possible values at each frame are the multiple possible detections of this keypoint. To obtain a cleaner model, we removed duplicate detections (within 7 pixels of each other) within each frame. To compensate for missed detected keypoints over many frames, we augmented the possible values at each frame with all detections up to *F* previous frames, weighted in time elapsed by multiplying their probability 2^−*F*^. We then identified the best path through the hidden Markov model using the Viterbi algorithm [95]. This procedure estimates a consistent path, even with missed detections of up to *F* frames.

#### Autoencoder filter

We found that the network would often try to predict a joint location even when the joint was occluded in that view. This type of error is particularly problematic when used in subsequent 3D triangulation. The convolutional neural network confidence scores associated with these predictions can be high, making them difficult to distinguish from correct, high-confidence predictions. To remove these errors, inspired by [96], we implemented a neural network that takes in a set of confidence scores from all keypoints in one frame, and outputs a corrected set of confidence scores. To generate a training set, we made use of the fact that human annotators do not label occluded joints but label all of the visible joints in each frame. Thus, we generated artificial scores from biased distributions to mimic what the convolutional neural network might predict for each frame, with visible joints given a higher probability on average. Specifically, we sample the scores from a normal distribution, with standard deviation of 0.3 and mean 0 for invisible and 1 for visible joints, clipped to be between 0 and 1. To mimic false positive or false negative detections, we flip 5% of the scores (*x* → 1 − *x*) at random. The task of the network is to predict a high score for each joint that is truly visible in that frame and a low score for any occluded joint. The network is a multilayer perceptron network with a single hidden layer and tanh activation units to perform this task. The size of the hidden layer is the number of joints (e.g. if there are 10 joint scores to predict, we set the hidden layer to 10 units). We trained the network using the Adam optimizer [97] implemented in the scikit-learn library [98]

### 4.4 Calibration of multiple cameras

#### Camera model

A camera captures 2D images of light reflecting from 3D objects; thus, we can think of each camera as a projection, transforming 3D vectors to 2D vectors. To establish our notation, for a point ***p*** = (*x, y, z*)^*T*^ or ***u*** = (*x, y*)^*T*^, we use a tilde to denote that point in homogeneous coordinates (with a 1 at the end), so that 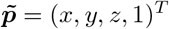 or 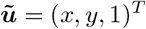.

A camera model specifies a transformation from a 3D point 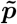 to a 2D point 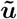. We use the camera model described by Zhang [64], which consists of a product of an intrinsics matrix **A**, an extrinsics matrix **P**, and a distortion function 𝒟.

The extrinsics matrix **P** ∈ ℝ^4×3^ describes how the camera is positioned relative to the world. We represent **P** as the product of a rotation matrix and a translation matrix. Both rotations and translations may be fully specified with 3 parameters each, for 6 parameters total in **P**.

The intrinsics matrix **A** ∈ R^3×3^ describes the internal coordinate system of the camera. It is often modeled using 5 parameters: focal length terms *f*_*x*_ and *f*_*y*_, offset terms *c*_*x*_ and *c*_*y*_, and a skew parameter *s*:

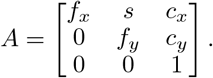

In practice, we found that we obtain a more robust calibration by reducing the number of parameters, setting *f* = *f*_*x*_ = *f*_*y*_, *s* = 0, and (*c*_*x*_, *c*_*y*_) to be at the center of the image, so that we need to estimate only the focal length parameter *f* for the intrinsics matrix.

The distortion function models nonlinear distortions in the camera pixel grid. This distortion is typically modeled with 3 parameters as

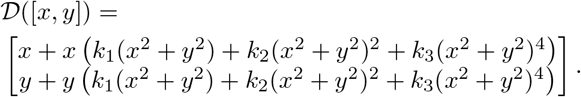

In practice, we found that the higher-order distortion terms *k*_2_ and *k*_3_ are often small for modern cameras, so we assume *k*_2_ = *k*_3_ = 0 and only estimate a single parameter *k*_1_.

Thus, the full mapping may be written as

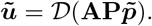

In total, the camera model involves estimating 8 parameters per camera: 6 for extrinsics, 1 for intrinsics, and 1 for distortion.

For the camera calibration and triangulation methods described below, we define the projection 𝒯 from 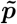 to 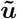 as

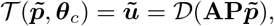

where ***θ***_*c*_ are the 8 parameters for the camera model of camera *c*.

#### Initial estimate of camera parameters

In order to calibrate the cameras and estimate parameters of the camera models, we start by obtaining an initial estimate of the camera parameters. We detected calibration board keypoints in videos simultaneously captured from all cameras. We then initialized intrinsics based on these detections following the algorithm from Zhang [64]. We initialized the distortion coefficients to zero.

We developed the following method to initialize camera extrinsics from arbitrary locations. For each pair of cameras, the number of frames in which the board is seen simultaneously is counted and used to build a graph of cameras. To be specific, each node is a camera, and edges represent pairs of cameras whose relation we will use to seed the initialization.

The greedy graph construction algorithm is as follows. Starting with the pair of cameras for which the number of frames the board is simultaneously detected is the largest, connect the two camera nodes with an edge. Next, proceed with iterations in decreasing order of the number of boards simultaneously detected. At each iteration, if the two nodes (cameras) are not already connected through some path, connect them with an edge. Processing iteratively through all pairs of cameras in this manner, a graph of camera connectivity is produced. Full 3D calibration is possible if and only if the graph is fully connected.

To initialize the extrinsics using this graph, we start with any camera and set its rotation and translation to zero. Then, we initialize its neighbors from the estimated relative pose of the calibration board between them using the initial intrinsics. This procedure is continued recursively until all cameras are initialized. A diagram of the camera initialization for an example dataset is provided in Figure 3A.

#### Bundle adjustment

To refine the camera parameters from initial estimates, we performed a bundle adjustment by implementing a nonlinear least-squares optimization to minimize the reprojection error [24]. Given all 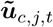, the detected *j*^th^ keypoints from the calibration board at cameras *c* in frames *t*, we solve for the best camera parameters ***θ***_*c*_ and 3D points 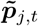 such that the reprojection loss ℒ is minimized:

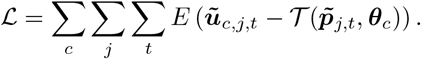

Here, *E*(·) denotes the norm using which the error is computed. This norm may be the least squares norm, but in practice, we used a robust norm, such as the Huber or soft *ℓ*_1_ norm, to minimize the influence of outliers.

This optimization is nonlinear because the camera projection function 𝒯 is nonlinear. We recognized that it is a nonlinear least-squares problem with a sparse Jacobian and thus solved it efficiently using the Trust Region Reflective algorithm [99, 100], as implemented in SciPy [101].

#### Iterative bundle adjustment

When calibrating cameras, we found that outliers have an outsized impact on calibration results, even when using robust losses such as the Huber loss or soft *ℓ*_1_ loss. Thus, we designed an iterative calibration algorithm, inspired by the fast global registration algorithm from Zhou et al. [102], which solves a minimization with a robust loss efficiently through an alternating optimization scheme.

We approximate this alternating optimization in the camera calibration setting through an iterative threshold scheme. In our algorithm, at each iteration, a reprojection error threshold is defined and the subset of points ***u***_*c,i*_ with reprojection error below this threshold is chosen. Bundle adjustment is then performed on these points alone. The threshold decreases exponentially with each iteration, to refine the points to be calibrated. The pseudocode for the algorithm is listed in Algorithm 1.

##### Algorithm 1 Iterative bundle adjustment

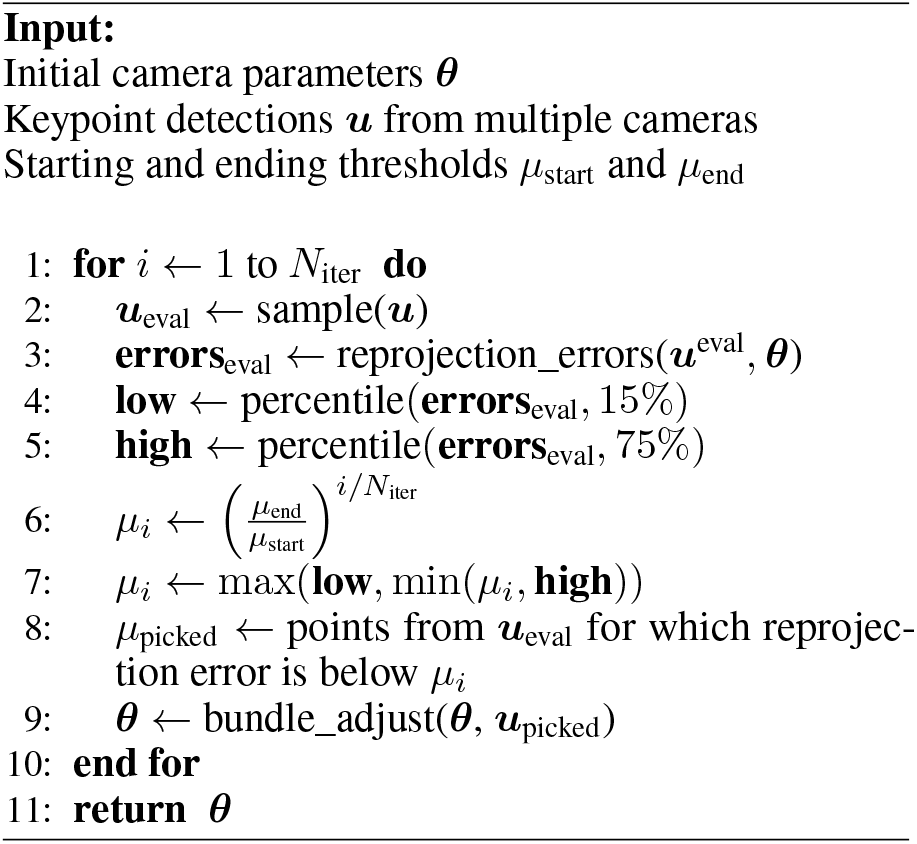

#### 4.5 Triangulation and 3D filtering

The 3D triangulation task seeks 3D points ***p***_*j,t*_ for joint *j* at frame *t*, given a set of detected 2D points ***u***_*c,j,t*_ from cameras *c* with camera parameters ***θ***_*c*_. There are several common methods for solving this triangulation task. Below, we describe 3 of these methods, then describe our method for spatiotemporally constrained triangulation. For illustration, a comparison of the performance of these methods is shown on an example dataset in Figure 7.

#### Linear least-squares triangulation

The first method triangulates 3D points by using linear least-squares [103]. Linear least-squares is the fastest method for multi-camera triangulation, but it may lead to poor results when the 2D inputs contain noisy or inaccurate keypoint detections. To be specific, we start with a camera model with parameters estimated from the calibration procedure described above, so that the extrinsics matrix **P**_*c*_, intrinsics matrix **A**_*c*_, and distortion function 𝒟_*c*_ are known for each camera *c*. By rearranging the camera model, we may write the following relationship:

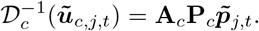

We solved this linear system of equations using the singular value decomposition (SVD) of the product **A**_*c*_**P**_*c*_ to approximate the solutions for the unknown 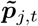 [103].

#### Median-filtered least-squares triangulation

As a simple extension of least-square triangulation to correct some of the noisy detections, we applied a median filter to the resulting 3D points tracked across frames. This filtering improves the tracking, but at the cost of losing high frequency dynamics. Furthermore, a median filter does not improve triangulation if the original tracking is consistently poor.

#### RANSAC triangulation

Random sample consensus (RANSAC) triangulation aims to reduce the influence of outlier 2D keypoint detections on the triangulated 3D point, by finding the subset of keypoint detections that minimizes the reprojection error. We implemented RANSAC triangulation by triangulating all possible pairs of keypoints detected from multiple views and picking the resulting 3D point with the smallest reprojection error.

Formally, let 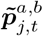 be the triangulated 3D point for keypoint *j* at frame *t* computed using the 2D keypoint detections from cameras *a* and *b*, then our algorithm finds 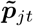 using the following relation:

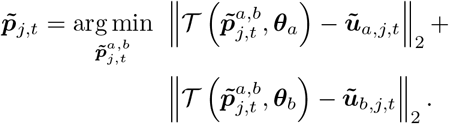

#### Spatiotemporally regularized triangulation

We formulated triangulation as an optimization problem, which allowed us to specify soft spatiotemporal constraints (i.e. regularization) on the triangulated points. We propose that the points must satisfy three soft constraints: (1) the projection of the 3D points onto each camera should be close to the tracked 2D points, (2) the 3D points should be smooth in time, and (3) the lengths of specified limbs in 3D should not vary too much. Each of these constraints may be formulated as a regularization in the full objective function.

First, the **reprojection loss** is written as

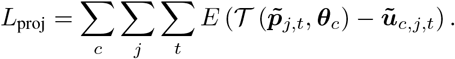

Here, *E*(·) is a robust norm function such as the Huber or soft-*ℓ*_1_ norm, to minimize the influence of outlier detections.

Second, the **temporal loss** is formulated as follows:

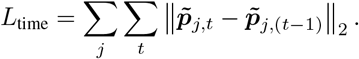

We extend this penalty to minimize higher-order (e.g. 2nd or 3rd) finite-difference derivatives, which produces smoother trajectories but has less impact on important high frequency dynamics (see Figure S5).

Third, the **limb loss** may be formulated by adding an additional parameter *d*_*l*_ for each limb *l*, defined to consist of joints *j*_1_ and *j*_2_:

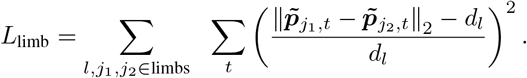

The limb error is normalized relative to the limb length so that each limb contributes equally to the error.

Given each of the losses above, the overall objective function to minimize may be written as:

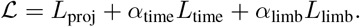

We solve this sparse nonlinear least-squares problem efficiently using the Trust Region Reflective algorithm [99, 100], as implemented in SciPy [101], similarly to the bundle adjustment optimization. To initialize the optimization, we use linear least-squares triangulation. When formulated as a sparse nonlinear least-squares problem, the time and memory requirements of the optimization scale linearly relative to the number of input time points.

The parameters *α*_time_ and *α*_limb_ may be tuned to adjust the strength of the temporal or limb loss, respectively. Note, however, that the temporal loss is in units of distance, which may vary substantially across datasets. Thus, to standardize these parameters, we break down the parameter *α*_time_ in terms of a user-tunable parameter *β*_time_ and an automatically computed scale *γ* such that

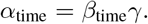

We compute the scale *γ* as

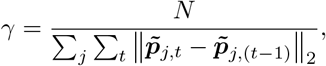

where 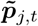 is an initial estimate obtained from linear least-squares triangulation. We found that the parameters *β*_time_ = 2 and *α*_limb_ = 2 work well across a variety of datasets, and we used these parameters for tracking all four datasets in this manuscript. The user may additionally specify weaker constraints for the lengths of certain limbs to allow for some flexibility, such as the shoulder length in humans or the length of the tarsus in flies.

#### Estimating joint angles

We estimated joint angles from the tracked 3D positions. To compute the flexion angle defined by the three 3D points surrounding the joint (***p***_*i*_, ***p***_*j*_, ***p***_*k*_), where point ***p***_*j*_ lies at the joint, the angle *f*_*j*_ is

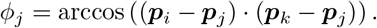

To estimate rotation and abduction angles, we solve an inverse kinematics problem treating the set of limb joints as a kinematic chain. When estimating limb angles from 3D coordinates of joints, the rotation of a joint is indistinguishable from the abduction of the next joint in the chain. We observed that fly and human limbs can be approximated to only have abduction at the joint closest to the body, so we resolve this ambiguity by assuming that only the first (most proximal) joint may abduct and the last (most distal) joint may not rotate.

The solution proceeds in two stages. In the first stage, we estimate the absolute rotation of each joint based on its {x, y, z} coordinate axes. The axes of the first joint match the coordinate system for the body. For other joints, the z axis is in the direction of the limb segment pointing from that joint away from the body, the x axis is in direction of proximal limb segment (towards the body) orthogonalized to the z-axis, and the y-axis is the cross product of the z-axis with the x-axis. In the second stage, the relative rotation between joints is computed and transformed to an Euler angle with an order of {z, y, x} for axis rotations. The rotations about the {z, y, x} axis represent rotation, flexion, and abduction angles, respectively. For more details of the implementation, see the accompanying code.

### 4.6 Evaluation

#### Comparison of bundle adjustment algorithms

To evaluate the different bundle adjustment algorithms (Figures 3B and S1), we ran the algorithms with different parameters on the calibration videos from the fly setup. There were 4475 frames where the calibration board was detected in 2 or more cameras. To demonstrate the usefulness of our iterative bundle adjustment procedure with lower number of detections, we evaluated all bundle adjustment algorithms after subsampling the frames with board detections to 313 (7%) and 4475 (100%). At each of these frame counts, we initialized the camera parameters and then ran our iterative bundle adjustment procedure, as well as traditional bundle adjustment with a linear least-squares loss, a Huber loss, and soft L1 loss. As the Huber and soft L1 losses are sensitive to the outlier threshold parameter, we evaluated them at multiple outlier thresholds on our dataset (Figure S1). We picked the loss with the best outlier threshold, as evaluated by the reprojection error at the 75th percentile, to plot in the main calibration figure. The iterative bundle adjustment procedure was run with the default parameters in Anipose: *N*_*iter*_ = 12, *µ*_start_ = 15, *µ*_end_ = 1.

#### Evaluation against physical ground truth

To evaluate the calibration and triangulation, we compared the accuracy of manual keypoint annotations, neural network keypoint detections, and OpenCV keypoint detections (Figure 4). The ground truth was considered to be known physical length and angles of the ChArUco board. The physical lengths were calculated between all pairs of keypoints by taking the length between the known positions of pairs of corners. Similarly, the physical angles were estimated between all triplets of non-collinear keypoints. The subpixel OpenCV detections were done using the Aruco module [25]. The manual annotation and neural network methods are detailed above. Given the keypoint detections from each method, we used linear least-squares triangulation to obtain 3D points and computed angles using the dot product method detailed above. If a keypoint was detected in fewer than 2 cameras at any time, we could not triangulate it and therefore did not estimate the error at that frame.

#### Evaluation of 3D tracking error for different filters

To evaluate the contribution of 2D and 3D filters, we applied each filter and measured the reduction in error. For the 2D filters, we applied each of the filters (2D median filter, Viterbi filter, and autoencoder filter) and computed the 3D position using linear least-squares triangulation. We could not train the autoencoder filter on the human dataset, as the filter relies on occluded keypoints not being present in the annotated dataset and, due to the nature of the human dataset, all keypoints are annotated from every view at every frame. When applying the spatiotemporal regularization, we assumed a low variance in length of the coxa, femur, and tibia in flies and of the arm, the forearm, pelvis, femur, and tibia in the human. We assumed a slightly higher variance for the length of the tarsus in each fly and of the neck and shoulders in each human, because these body segments are more flexible. The parameters for each filter are listed in Table S1. We measured the error in joint positions and angles relative to those computed from manual annotations, using the *ℓ*_2_ norm. To evaluate the effect of the filter addition, as there was a lot of variance in error across points, we computed the difference in error for each point tracked. We treated points with reprojection error above 20 pixels as missing. The procedure for evaluating the 3D filters was similar, except that we compared the error in joint position and angle relative to the error from 3D points obtained with a Viterbi filter and autoencoder filter with linear least-squares triangulation.

#### Evaluation of derivative error for different filters

To evaluate the contribution of different 2D and 3D filters to the error in derivative estimation, we applied each filter to the 3D trajectory of each joint and estimated the derivative by using the finite difference method. For each joint, each frame, and each filter, we obtain a 3D vector representing a derivative. We compare the error between this derivative vector and the true derivative vector from manual annotations by using the *ℓ*_2_ norm, as in the previous section.

#### Evaluation of 3D tracking error for different number of cameras

To evaluate how the number of cameras contributes to the estimate of error, we ran Anipose on all combinations of 2, 3, and 4 cameras for the human dataset. We measured the error in joint position and angles relative to manual annotations as described above. We plotted the mean error across all joint positions or angles and across all possible combinations of cameras (Figure S6) at each number of cameras.

#### Evaluation of temporal regularization on synthetic dataset

To evaluate how minimizing higher order derivatives affects tracking of high frequency movement dynamics, we evaluated the temporal regularization on a synthetic dataset (Figure S5). We synthesized 30 ground-truth keypoint trajectories, each of length 500, by applying a low-pass filter with a cutoff of 0.12 cycles/sample on white noise. We then corrupted these trajectories by adding white noise and removing 10% of the points, simulating observed triangulated points (for example, as in the “No filters” trace in Figure 7A). We reconstructed the signal using temporal regularization and minimizing the 1st, 2nd, or 3rd derivative across different levels of smoothing factor *β*_time_. We estimated the power spectrum of the ground truth, corrupted, and reconstructed signals by taking the average power spectral density at each frequency across all 30 simulated trajectories. We estimated the power spectral density using the Welch’s method as implemented in SciPy [101]. We computed the root mean squared error (RMSE) between the ground truth and reconstructed signals for each derivative minimized at different levels of smoothing. We evaluated the RMSE of median filters with window size of 3 to 25 samples on the same trajectories, and found the median filter with a window size of 9 samples to have the lowest RMSE, which we plot as a reference.

### 4.7 Analysis of kinematics

#### Analysis of fly walking kinematics

For the analysis in Figure 10, we used data from 39 wild-type Berlin flies on a spherical treadmill (details of experimental setup above). We tracked the flies using Anipose with spatiotemporal regularization and Viterbi and autoencoder filters. We confirmed by visual inspection and by checking reprojection errors that all flies were well tracked.

To restrict the data to only walking, we manually labeled fly behavior for a random subset of videos using the VGG Image Annotation tool [91]. The categories of behaviors labeled were abdomen grooming, antennae grooming, ball push, ball tapping, eye grooming, head grooming, standing, t1 grooming, t3 grooming, walking. To detect walking behavior across the entire dataset, we fit a logistic classifier to predict the type of behavior. The input data to the classifier for each time point was a chunk of 24 samples around that time of 3D joint positions and angles and the Fourier transform of the 24 samples of each variable. The confusion matrix for the classifier on a test set is shown in Figure S8C. The false negative rate was 0%, whereas the false positive rate was about 3%. To detect bouts of walking, we used the classifier to predict a walking probability for each sample in a video, applied a mean filter with a window of 16 samples to the probability, then kept bouts where the probability was above 0.5 for at least 40 consecutive samples. To further reduce spurious walking bout detections, we removed any bout where the femur-tibia flexion of the left front and hind legs varied less than 10 degrees over the full bout. We confirmed with visual inspection that all bouts removed in this way did not include walking.

To perform the UMAP embeddings, we followed a procedure inspired by DeAngelis et al [27], which mapped the manifold structure of *Drosophila* walking from 2D tracking data. We took chunks of 32 samples, advancing by 8 samples, of the coxa rotation, femur rotation, and femur-tibia flexion angles and their derivatives. Thus, we obtained a set of vectors of size 1152 (32 samples * 6 legs * 3 angles * 2 raw & derivatives), which we standardized by subtracting the mean and dividing by the standard deviation along each dimension. We embedded this set of vectors in 3 dimensions using the UMAP algorithm [31], with effective minimum distance of 0.4 and 30 neighbors as parameters. To compute the phase of the step cycle, we applied a band-pass filter (1st order Butterworth over 3– 60Hz) to front left leg femur-tibia flexion and estimated the phase from the analytic signal obtained using the Hilbert transform.

#### Analysis of mouse reaching kinematics

In Figure S9, we analyzed videos from 4 mice recorded over 2 different days (details of experimental setup above). We tracked 3 keypoints on the hand for each mouse using Anipose with no filters. To obtain accurate 3D tracking for all trajectories, we removed all points with reprojection error above 10 pixels, then filled in missing data (about 11% of the data) using linear interpolation. We used the proximal end of digit 3 as a marker for the overall hand position. Mice 1 and 3 reached with their left hand, whereas mice 2 and 4 reached with their right hand. Accordingly, we quantified the movement of the hand each mouse reached with. We labeled the start and end of each reach, along with the reach type using the Anipose visualizer (Figure 9). To obtain the 3D position of the pellet holder, we labeled the pellet holder for each mouse and day from both views using the VGG Image Annotation tool [91], then triangulated the labeled points for each pair of views using aniposelib. We measured the distance of the hand (proximal end of digit 3) to the pellet holder by using the *ℓ*_2_ norm.

#### Analysis of human walking kinematics

In Figure S10, we analyzed videos from all 7 publicly available subjects in the Human 3.6M dataset (dataset described above). We tracked 17 keypoints for each human using Anipose with spatiotemporal regularization and Viterbi filters.

To focus on walking, we restricted our analysis on the “Walking-1”, “Walking-2”, “WalkingTogether-1”, and “WalkingTogether-2” actions in the dataset. We estimated the knee flexion, hip flexion, and hip rotation angles as described in the “Estimating joint angles” section above. For the UMAP embedding, we followed a procedure similar to our analysis of fly kinematics. Specifically, we took chunks of 24 samples, advancing by 8 samples, of the knee flexion, hip rotation, and hip flexion angles and their derivatives. Thus, we obtained a set of vectors of size 288 (24 samples * 2 legs * 3 angles * 2 raw & derivatives), which we standardized by subtracting the mean and dividing by the standard deviation along each dimension. We embedded this set of vectors in 3 dimensions using the UMAP algorithm [31], with effective minimum distance of 0.4 and 30 neighbors as parameters.

## Contributions

PK, BWB, and JCT conceived the project. PK designed, implemented, and evaluated the Anipose toolkit. KR wrote the Anipose documentation, contributed Tensorpack data augmentation to DeepLabCut, and wrote key parts of the Anipose visualizer. ESD and SWB collected the ChArUco and fly datasets. ES and EA collected the mouse dataset. PK, BWB, and JCT wrote the paper, with input from KR, ESD, ES, and EA.

## Acknowledgments

We thank Su-Yee Lee and Chris Dallmann for help in annotating keypoints on flies, John So for help with annotating keypoints on the ChArUco board, and Sam Mak for help with annotating mice keypoints and fly behavior. We thank Stephen Huston for loan of his calibration board and Julian Pitney for contributing code to check calibration board detections to Anipose. Finally, we thank the Tuthill and Brunton labs and Mackenzie and Alexander Mathis for support, suggestions, and feedback on the manuscript.

PK was supported by a National Science Foundation Graduate Research Fellowship. KLR was supported by fellowships from the University of Washington’s Institute for Neuroengineering (UWIN) and Center for Neurotechnology (CNT). ESD was supported by a fellowship from University of Washington’s Institute for Neuroengineering. ES was supported by the National Institutes of Health (F31NS115477). EA was supported by the National Institutes of Health (R00 NS088193, DP2NS105555, R01NS111479, and U19NS112959), the Searle Scholars Program, The Pew Charitable Trusts, and the McKnight Foundation. BWB was supported by a Sloan Research Fellowship and the Washington Research Foundation. JCT was supported by the Searle Scholar Program, the Pew Biomedical Scholar Program, the McKnight Foundation, and National Institute of Health grants R01NS102333 and U19NS104655. JCT is a New York Stem Cell Foundation – Robertson Investigator.

## Supplemental figures

**Table S1:** Anipose configuration parameters used in this paper. Related to Figures 6, 7, 8.

|  | ChArUco | Mouse | Fly | Human |
| --- | --- | --- | --- | --- |
| Training frames | 1200 | 2200 | 6632 | 636724 |
| Test frames | 1200 | 400 | 1200 | 159181 |
| Num cameras | 6 | 2 | 6 | 4 |
| Pixel scale (mm) | 0.0075 | 0.0897 | 0.0075 | 4.79 |
| **2D filter** |  |  |  |  |
| score threshold |  |  | 0.05 | 0.05 |
| n_back |  |  | 3 | 3 |
| medfilt |  |  | 13 | 13 |
| offset_threshold |  |  | 15 | 30 |
| spline |  |  | true | true |
| **3D filter** |  |  |  |  |
| score_threshold | 0.3 | 0.3 | 0.3 | 0.3 |
| reproj_error_threshold |  |  | 5 | 5 |
| scale_length |  |  | 3 | 1.5 |
| scale_length_weak |  |  | 0.5 | 0.5 |
| scale_smooth |  |  | 2 | 4 |
| n_deriv_smooth |  |  | 3 | 2 |

**Figure S1:**
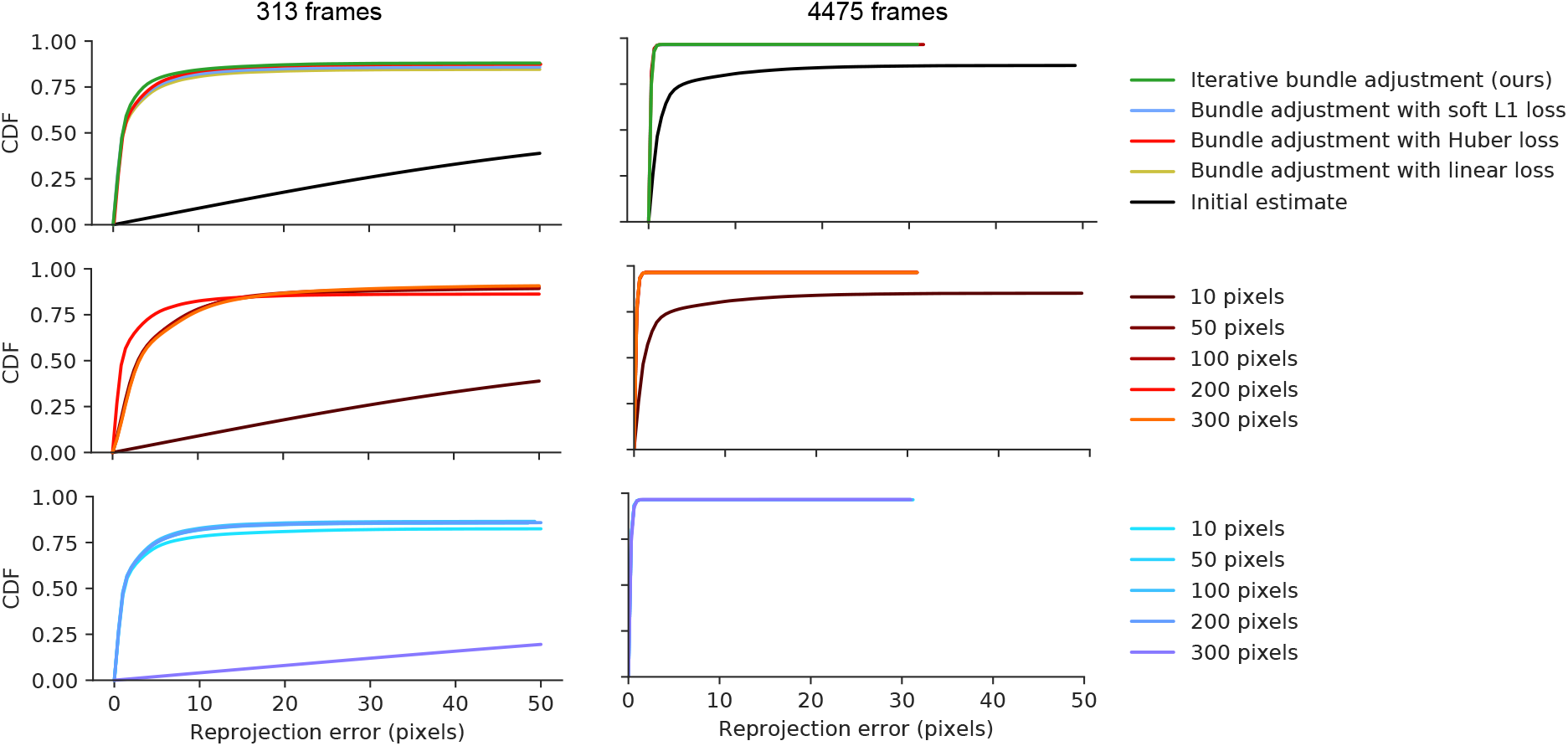
Related to Figure 3. Reprojection error as a function of outlier threshold for bundle adjustment with Huber and soft L1 losses.

**Figure S2:**
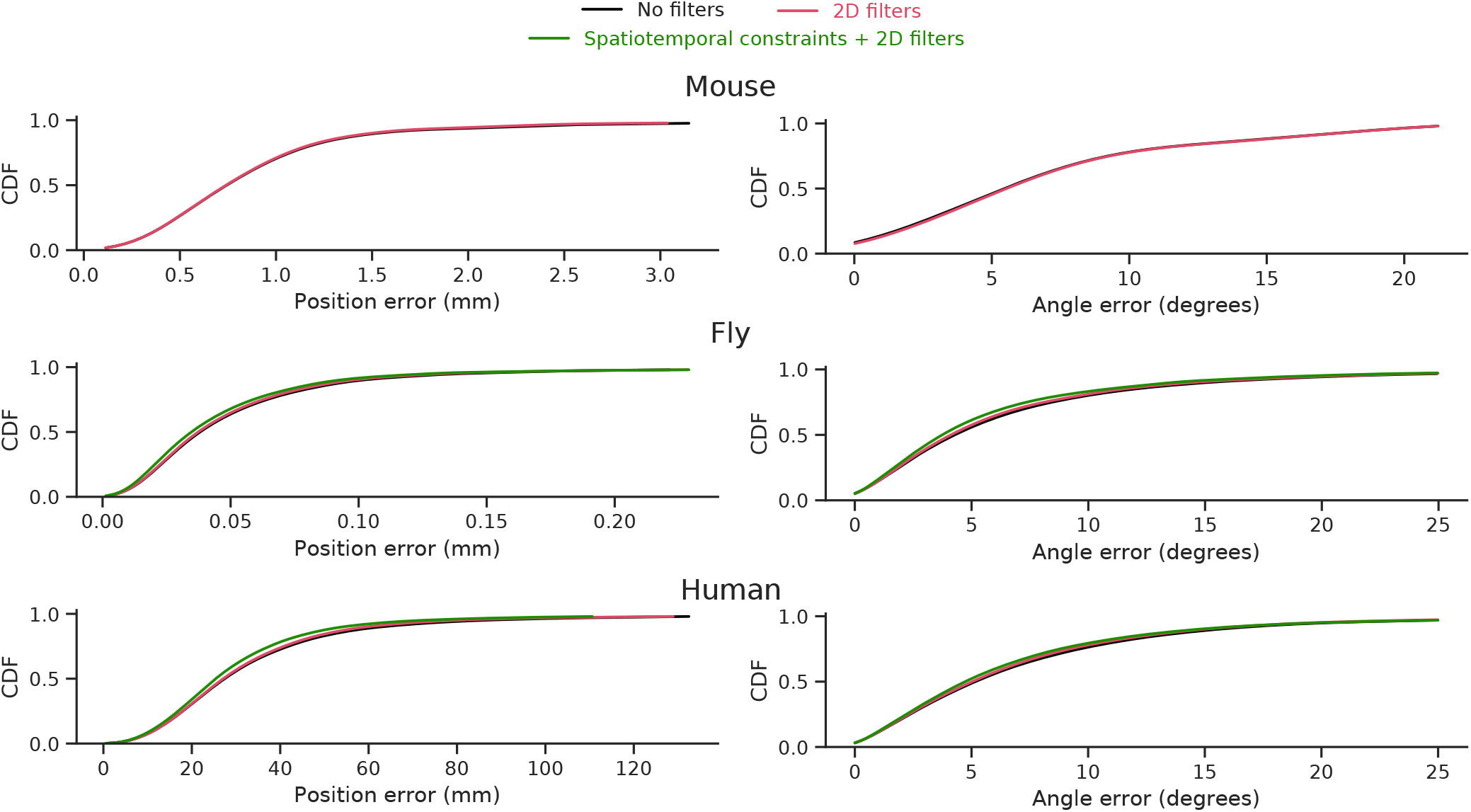
Cumulative distribution functions of the position and angle error with and without filters for each of the datasets. Related to Figure 5

**Figure S3:**
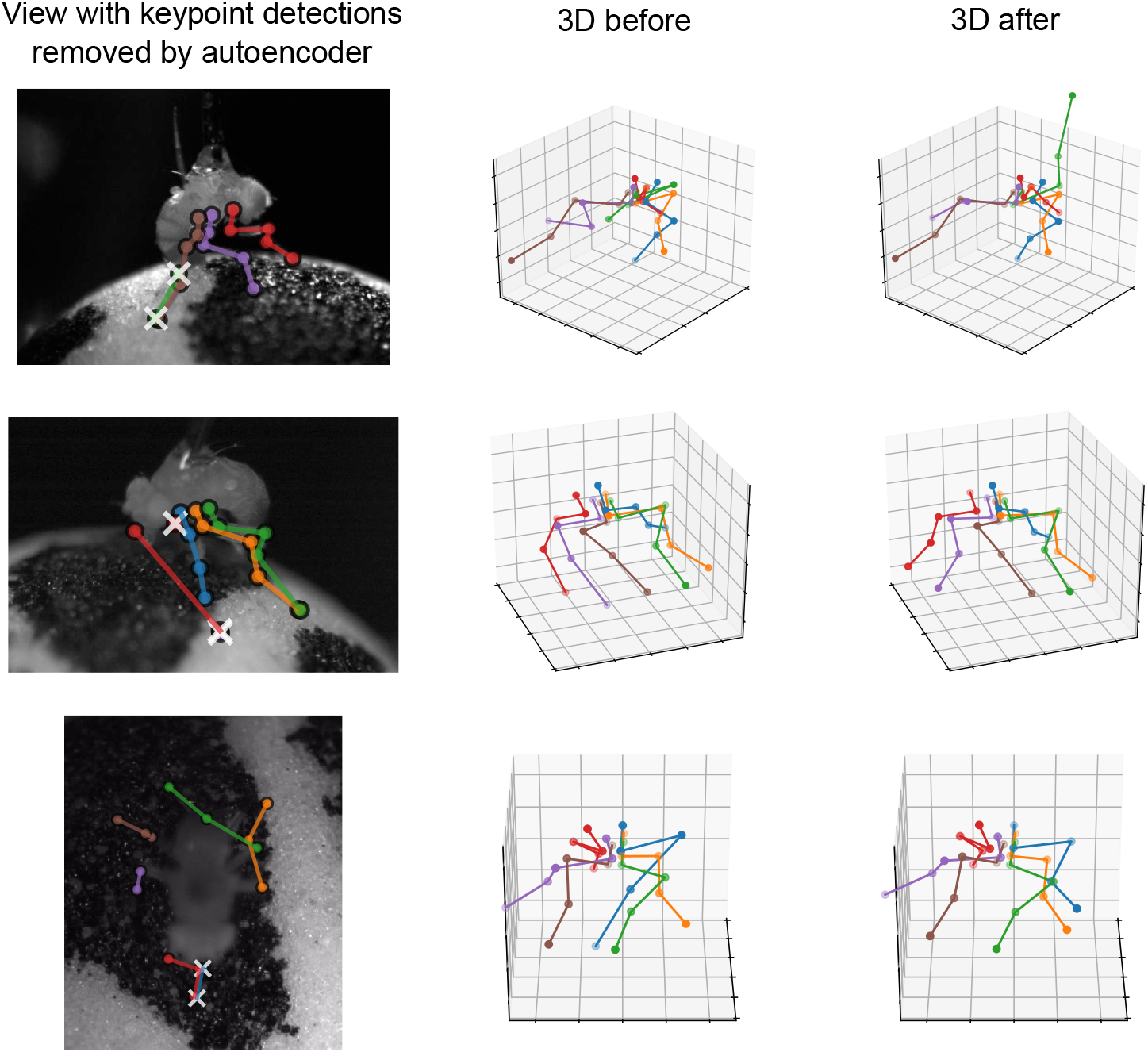
An autoencoder corrects 3D tracking by removing bad keypoint detections. Related to Figure 6. On the left is one view where the autoencoder lowered the confidences for particularly bad detections, thus removing them from the 3D triangulation. On the right are the 3D positions of the keypoints before and after the removal.

**Figure S4:**
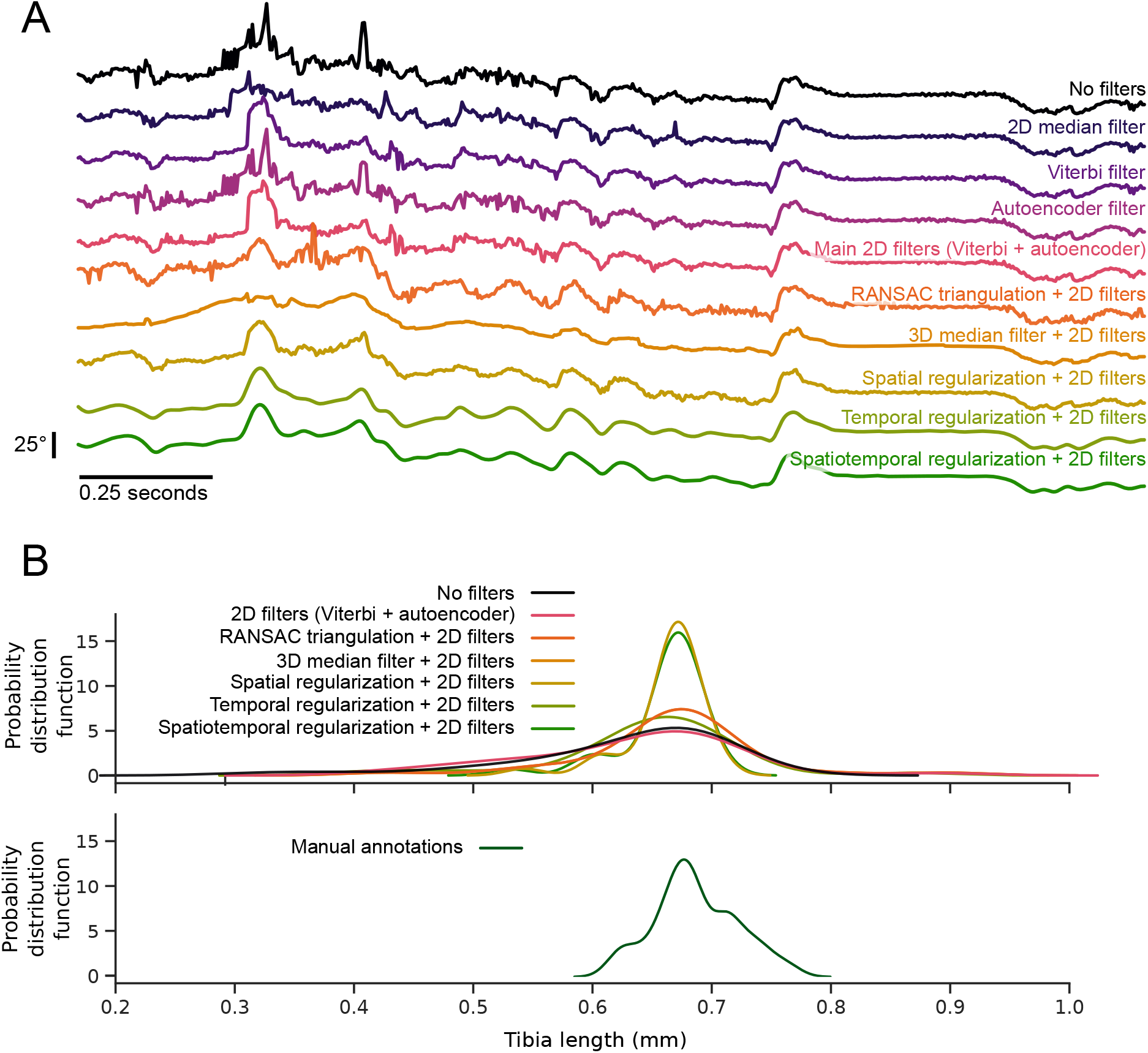
Related to Figure 7. (A) Example traces of the tracked hind-leg femur tibia flexion angle, before and after filtering. (B) Comparison of methods for estimating tibia length. Spatial regularization most closely matches the distribution of tibia lengths based on manual annotations. The plots show the distribution of tibia lengths for one fly, extending the example shown in Figure 7B, for different filtering strategies (top) and manual annotations (bottom).

**Figure S5:**
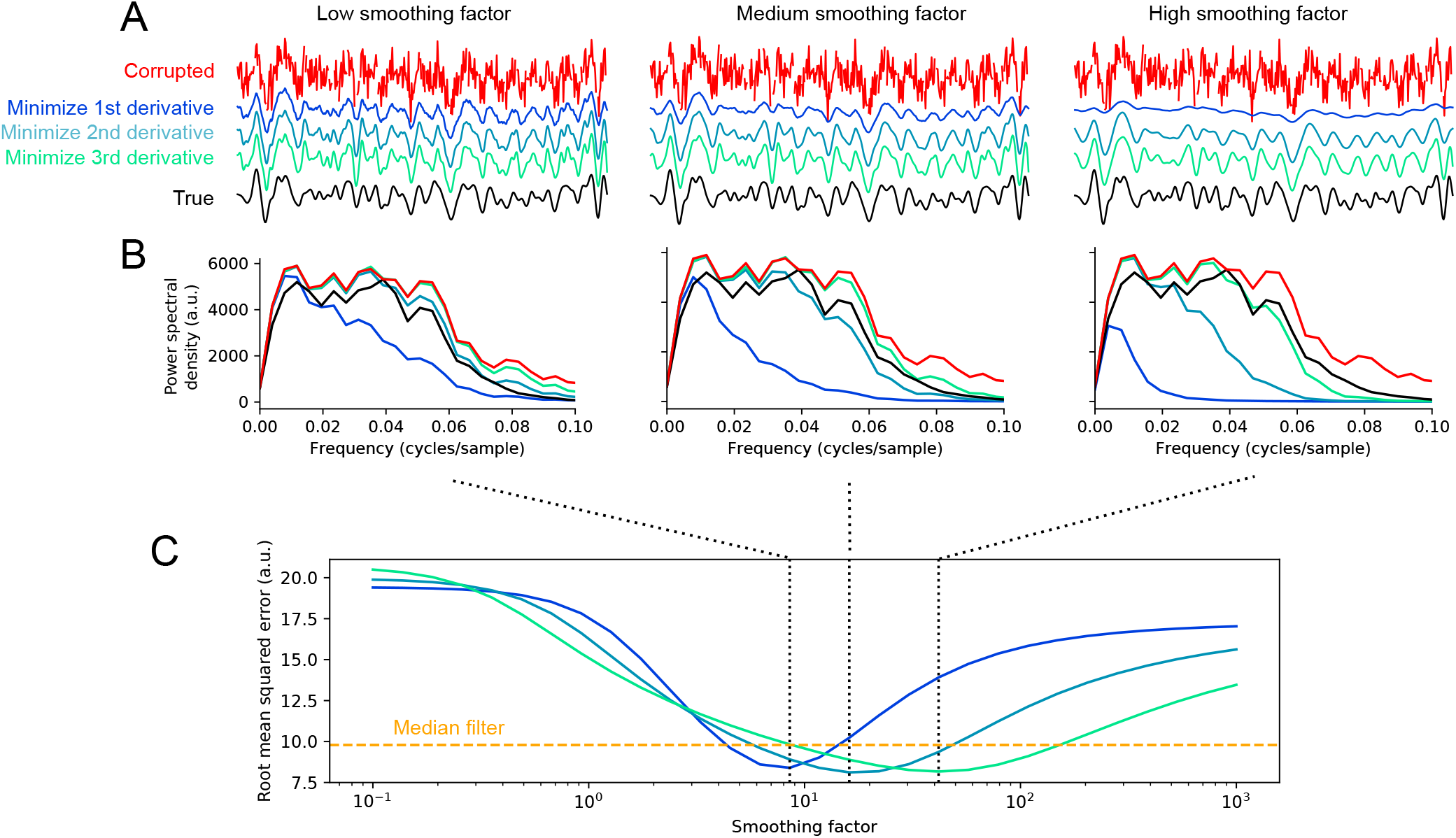
Minimizing higher order derivatives preserves high frequency dynamics and leads to lower reconstruction error. Related to Figure 7. (A) An example simulated trajectory along with its reconstructions using temporal regularization with different derivatives minimized. Each column shows reconstructions with different smoothing factors. (B) We synthesized 30 different trajectories with the procedure in A and compared the average power spectral density between the true, corrupted, and reconstructed trajectories with different derivatives minimized. At any smoothing factor, minimizing higher derivatives preserves more power at high frequencies. (C) The average root-mean squared error (RMSE) of reconstruction for the 30 simulated trajectories. The minimum error for a median filter (over all possible filter widths) is shown as a dashed line, for reference. Dotted lines indicate the smoothing factors shown in A and B. Note that minimizing higher derivatives is more robust to smoothing factor choice, as a wider range of factors give lower RMSE than a median filter. The best RMSE over all possible smoothing factors is lower when minimizing the 3rd derivative than 2nd or 1st.

**Figure S6:**
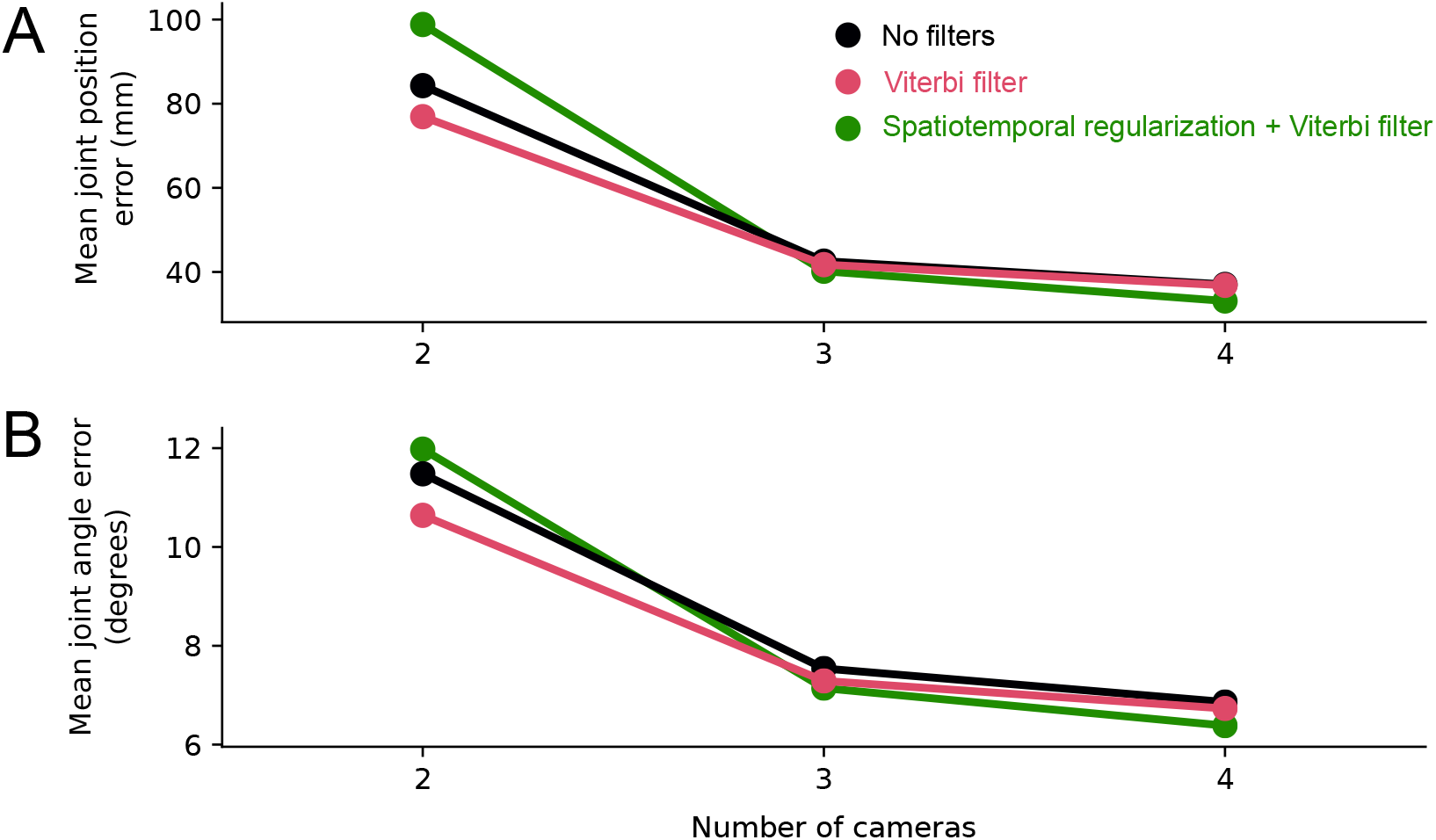
Estimates of error with different number of cameras for the human dataset. Related to Figure 7. (A) Mean joint position error of Anipose tracking with subsets of cameras and different filter conditions. (B) Same as A, but plotting mean joint angle error.

**Figure S7:**
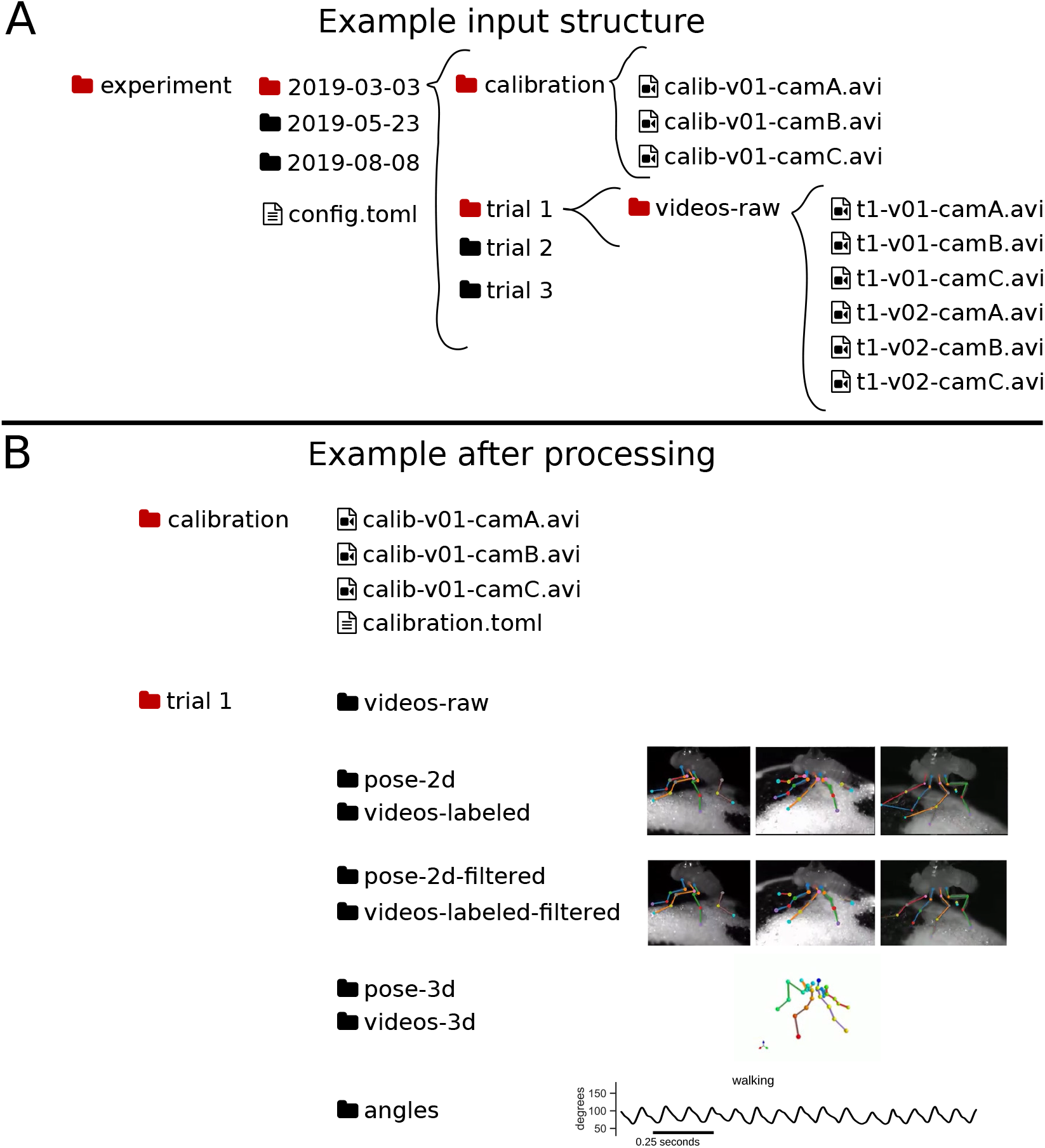
An example of the Anipose file structure. This structure enables visualization of arbitrary datasets, as shown in Figure 9. (A) The input file structure consists of folders nested to arbitrary depths (e.g. “experiment/2019-03-03/trial 1”) with a folder for raw videos at each leaf of the directory tree. The calibration folder may be placed anywhere and will apply recursively to all folders adjacent to it. (B) When the user runs Anipose, it will create a folder for each step of processing. New folders created include “pose-2d” and “videos-labeled” which contain the unfiltered keypoint detections and visualizations of those, “pose-2d-filtered” and “videos-labeled-filtered” which contain the filtered keypoint detections and visualizations, “pose-3d” and “videos-3d” which contain the triangulated 3D keypoint detections and visualizations of these, and finally “angles” which contains angles computed based on the 3D keypoint detections.

**Figure S8:**
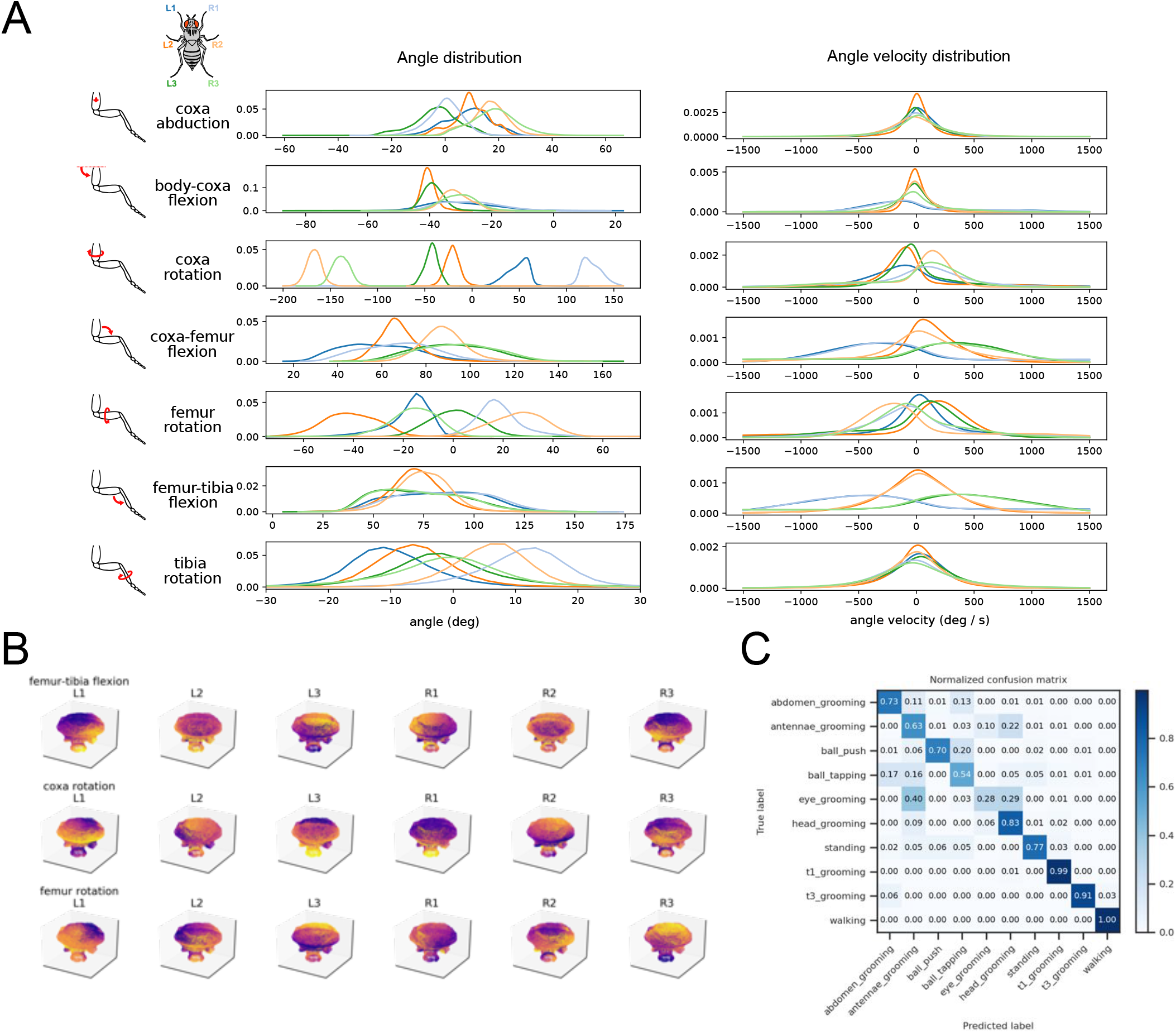
Related to Figure 10. (A) Probability density functions of all joint angles and derivatives for 39 wild type flies during walking, extending the subset of angles presented in Figure 10B. (B) UMAP embeddding of fly walking, as in Figure 10C, colored by each of the joint angles. The colormap is normalized to the angle within each plot. (C) Confusion matrix for the behavior classifier used to isolate walking bouts for Figure 10.

**Figure S9:**
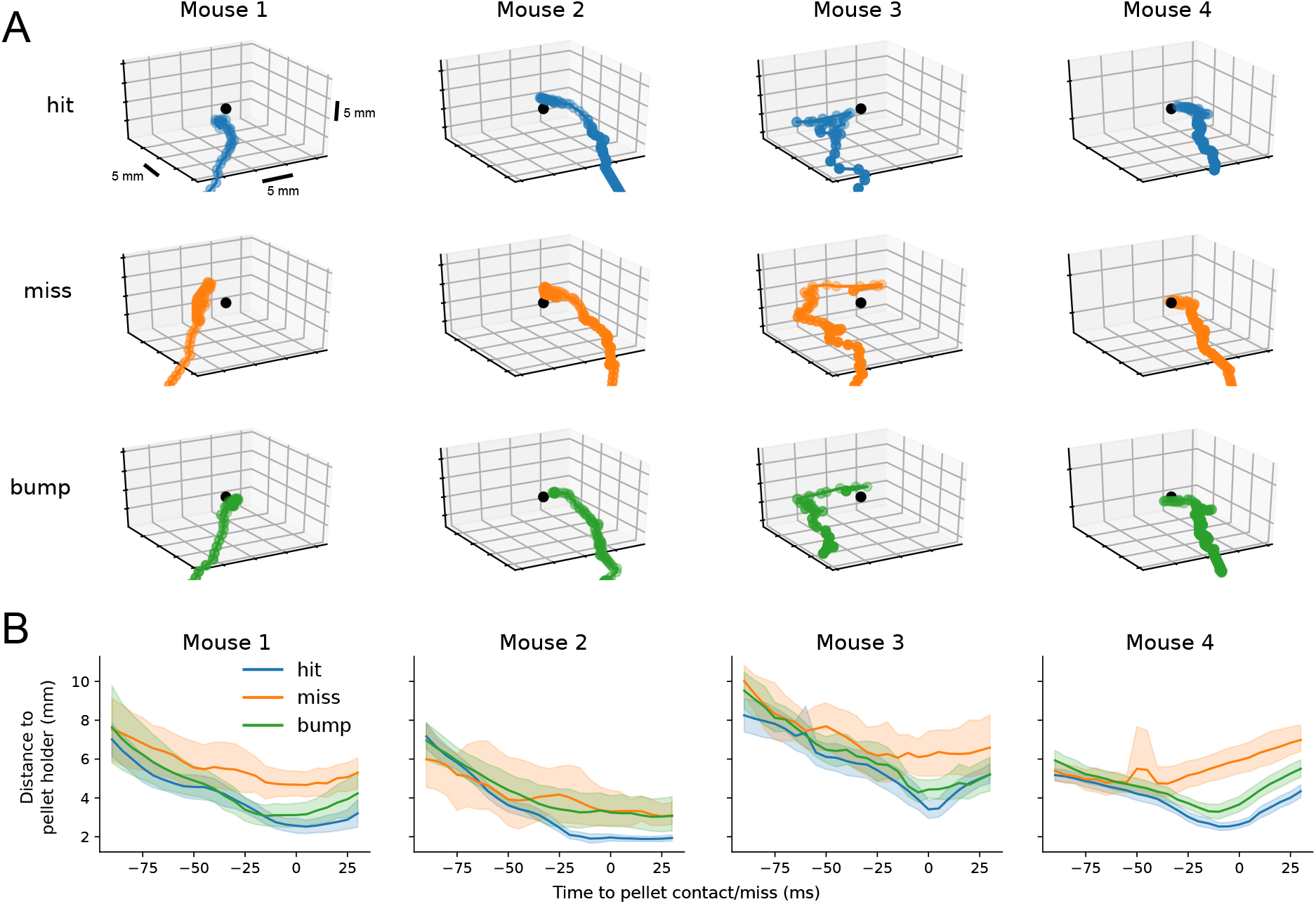
3D tracking with Anipose reveals common structure of mouse reaches. (A) 3D trajectories of example reaches of each type. The pellet holder is indicated as a black dot. (B) Mean distance to pellet holder as a function of time, for each mouse. Shaded areas are 95% confidence intervals. When reaches are aligned to grasp attempt (0 ms), the hand is farther from the pellet on miss trials compared to hit or bump trials.

**Figure S10:**
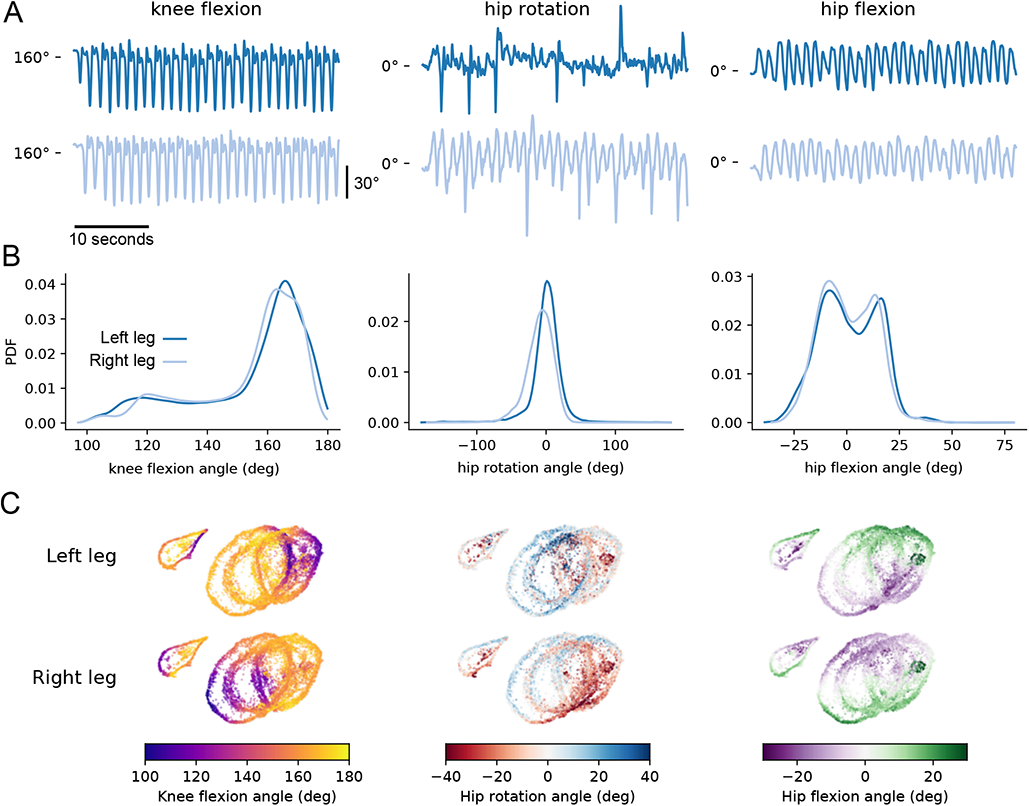
3D tracking of human walking enables quantification of leg angles and comparison across individuals. (A) Representative traces of knee flexion, hip rotation, and hip flexion from a walking human, tracked with Anipose. Data is from the Human 3.6M dataset. The median angle value is indicated at left as a reference point. (B) Probability distribution functions of knee flexion, hip rotation, and hip flexion angles from 7 humans. Only sessions that include walking are included. Note the asymmetry in the distributions of knee flexion and hip flexion, revealing the known non-sinusoidal pattern of knee and hip flexion during walking. (C) Axis units are arbitrary. Although each human subject has a characteristic gait, there is a continuum across all subjects. (D) UMAP embedding of knee flexion, hip rotation, and hip flexion angles across all legs, and their derivatives. The UMAP embedding is colored by knee flexion and hip rotation for each leg. Coloring by knee flexion angle reveals the common phase alignment of the circles across subjects. From this phase alignment, we see that the trajectory of hip rotation for each subject is markedly different.

